# Selective sweep probabilities in spatially expanding populations

**DOI:** 10.1101/2023.11.27.568915

**Authors:** Alexander Stein, Ramanarayanan Kizhuttil, Maciej Bak, Robert Noble

## Abstract

Evolution during range expansions shapes biological systems from microbial communities and tumours up to invasive species. A fundamental question is whether, when a beneficial mutation arises during a range expansion, it will evade clonal interference and sweep through the population to fixation. However, most theoretical investigations of range expansions have been confined to regimes in which selective sweeps are effectively impossible, while studies of selective sweeps have either assumed constant population size or have ignored spatial structure. Here we use mathematical modelling and analysis to investigate selective sweep probabilities in the alternative yet biologically relevant scenario in which mutants can outcompete and displace a slowly spreading wildtype. Assuming constant radial expansion speed, we derive probability distributions for the arrival time and location of the first surviving mutant and hence find surprisingly simple approximate and exact expressions for selective sweep probabilities in one, two and three dimensions, which are independent of mutation rate. Namely, the selective sweep probability is approximately (1 *− c*_wt_/*c*_m_)^*d*^, where *c*_wt_ and *c*_m_ are the wildtype and mutant radial expansion speeds, and *d* the spatial dimension. Using agent-based simulations, we show that our analytical results accurately predict selective sweep frequencies in the two-dimensional spatial Moran process. We further compare our results with those obtained for alternative growth laws. Parameterizing our model for human tumours, we find that selective sweeps are predicted to be rare except during very early solid tumour growth, thus providing a general, pan-cancer explanation for findings from recent sequencing studies.

## INTRODUCTION

Range expansion – the spatial spread of populations into new regions – is ubiquitous across biological scales and alters the course of evolution in distinct, often profound ways that remain incompletely understood [1]. Among cell populations, evolution during range expansions determines the development and spatial heterogeneity of biofilms [2], tumours [3], mosaicism [4], and normal tissue [5]. At the species level, range expansions influenced human evolution [6] and are of growing importance as climate change forces organisms into new habitats [7, 8]. Prior theoretical and experimental investigations of evolution during range expansion have considered the case in which the wildtype population spreads into new territory much faster than any mutant can displace the wildtype (for example, [9–12]). In this scenario, which is typical of microbial colonies growing *in vitro*, mutants can expand only at the population boundary and selective sweeps are precluded. The alternative case of slow range expansion and strong selection has been unexplored but is widely plausible. Consider, for example, an invasive species that is rapidly adapting to its new environment while gradually displacing a resident competitor, or a bacterial colony whose growth is slowed by sub-inhibitory antibiotic treatment.

Somatic evolution provides especially strong motivation for investigating the likelihood of selective sweeps during slow range expansions. Cancer results from an accumulation of mutations that drive cells to proliferate uncontrollably and to invade surrounding tissue [13]. An early, highly influential model based on colorectal cancer genetics posited a linear mode of evolution in which tumours acquire mutations via sequential selective sweeps [14]. Yet recent advances in multi-region and single-cell sequencing have revealed pervasive genetic intratumour heterogeneity [15, 16]. Although fitter mutant clones arise throughout cancer progression, they apparently seldom achieve selective sweeps, except perhaps in very small tumours [17]. These observations have sparked an ongoing debate about the general nature of tumour evolution, the extent of clonal selection, and the causes of intratumour heterogeneity [18, 19]. Mathematical modelling offers a way to resolve the conflict but, with notable exceptions [20–22], theoretical investigations of cancer initiation and early evolution have either ignored spatial structure [23–25] or have relied on agent-based models [26–29]. The typical observed pattern of very early fixation events followed by branching or effectively neutral intratumour evolution [18, 19] – which has important clinical implications [30–32] – has eluded a general explanation.

Here we use mathematical analysis to explain why beneficial mutations typically fixate only – if at all – in the very early stages of range expansions, even when mutants can displace the wildtype faster than the wildtype expands its range. We focus on a simple model with short-range dispersal throughout the population. By solving our model in the canonical case of constant radial expansion speed we derive exact and simple approximate expressions for sweep probabilities in one, two and three dimensions. We confirm the accuracy and robustness of our analytical results using extensive agent-based simulations of a spatial Moran process, and we compare outcomes for alternative growth models. We discuss how these findings shed light on the nature of evolution in range expansions in general, and cancer development in particular.

## RESULTS

### Macroscopic model

We consider a wildtype population that starts expanding spherically at time *t* = 0, such that its radius *x*_wt_ grows at speed *c*_wt_. Focusing on selective sweeps, we consider only advantageous mutations, which we assume spread within the wildtype at speed *c*_m_ > *c*_wt_ (Figure 1). Mutations occur at per-capita rate 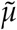 with each surviving genetic drift with probability *ρ*. In our analytic model, it is enough to consider the compound parameter 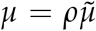, which is the mutation rate conditioned on survival. For brevity, we will refer to *μ* as the mutation rate unless otherwise mentioned.

**Figure 1.**
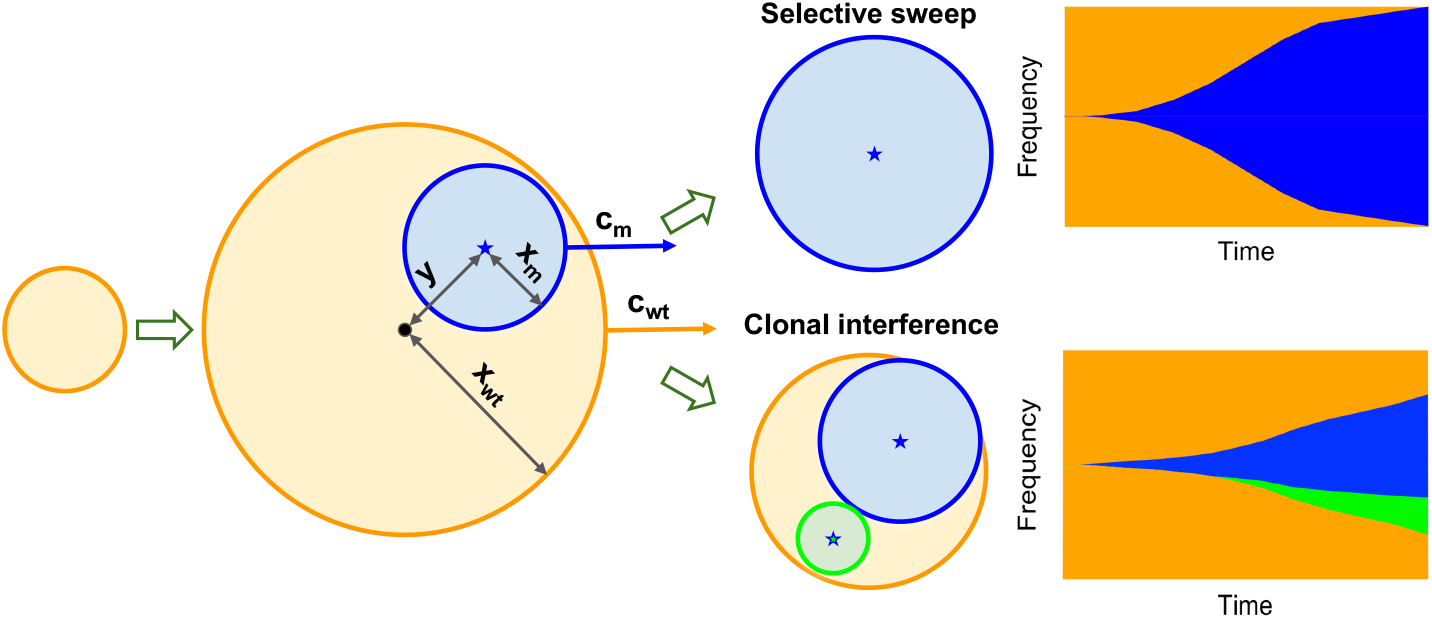
Illustration of the macroscopic model (not to scale) showing the two possible fates of the first surviving mutant (blue) within the wildtype population (orange). In the first case, the mutant reaches every part of the wildtype boundary before further mutants arise in the wildtype, and so achieves a selective sweep. Otherwise, a second (green) mutant arises in the wildtype population and generates clonal interference.

For mathematical and biological reasons (see Discussion), we focus on a model with constant radial expansion speeds; later we compare these results with those that pertain for alternative growth models. Various models link propagation speeds to fitness values and migration rates. The most prominent formula is obtained from a reaction-diffusion equation associated with Fisher [33] and Kolmogorov [34]: 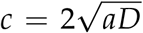, where *D* is the diffusion coefficient carrying information of migration and *a* is the difference in the local proliferation rate. More recent studies have sought to refine and generalise this result [35–38]. Our main results apply to any model that generates approximately constant speeds.

The first surviving mutation achieves a complete (or “hard” or “classic”) selective sweep only if it reaches every part of the wildtype expansion front before a second mutant of equal or greater fitness arises within the wildtype (Figure 1). Otherwise, the outcome is clonal interference or possibly a multiple-origin soft selective sweep if the competing mutations are sufficiently similar [39]. For simplicity, we neglect mutants with fitness values between those of the wildtype and the first mutant, which would slow the expansion of the first mutant and so reduce rather than negate the selective sweep probability. Neither do we investigate the case of a yet-fitter mutant that arises from the first mutant and achieves a selective sweep.

### Sweep probability

The unconditional sweep probability is derived in four steps. First, we introduce the random variable *X*, the radius of the wildtype population when the first mutant arises, and we compute its probability density *f*_*X*_(*x*). Second, we introduce the random variable *Y*, the distance between the wildtype and mutant origins, and we calculate its probability density conditioned on *X*, namely *f*_*Y*_(*y*|*X* = *x*). Third, we derive an expression for the conditional sweep probability Pr(sweep|*X* = *x, Y* = *y*). Finally, we marginalize out *X* and *Y* to obtain the unconditional sweep probability Pr(sweep). Here, we focus on the three-dimensional case; analogous results in one and two dimensions are presented in the SI Text.

#### Probability that no mutations occur

The following result (proved in SI Text) will be useful in various parts of subsequent derivations.

**Claim 1**. *The probability that k mutations arise and survive during the time interval* [0, *t*] *is Poisson distributed*,

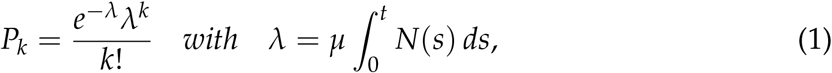

where *N*(*t*) *is the population size. In particular, the probability that no successful mutant arises is P*_0_ = *e*^−λ^.

Although it is commonly assumed that mutations are coupled to divisions, it is straightforward to translate between per-capita and per-division mutation rates (see SI text).

#### Arrival time of the first mutant

In the absence of mutants, the wildtype population in three dimensions grows as 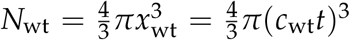. Applying Claim 1, we obtain the probability that no mutations arise in the time interval [0, *t*],

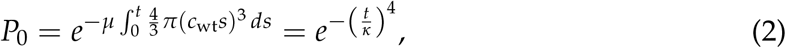

in terms of a characteristic duration 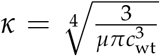. We identify 1 *− P*_0_ as the cumulative distribution function for the arrival time *T* of the first surviving mutant. The probability density of *T* is then

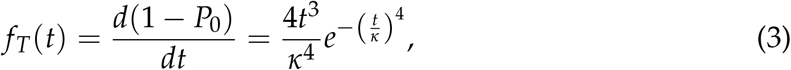

which is the Weibull distribution with shape parameter 4 and scale parameter *κ*. Substituting *x* = *c*_wt_*t*, we find that *f*_*X*_(*x*), the probability density of the radius of the wildtype population *X* at the time the first surviving mutant arises, is the Weibull distribution with shape parameter 4 and scale parameter 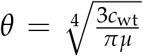. It follows that E[*X*] *≈* 0.91 *θ* and Var[*X*] *≈* 0.065 *θ*^2^. Analogous calculations in one and two dimensions yield similar Weibull distributions (SI text; Figure 2A-C).

**Figure 2.**
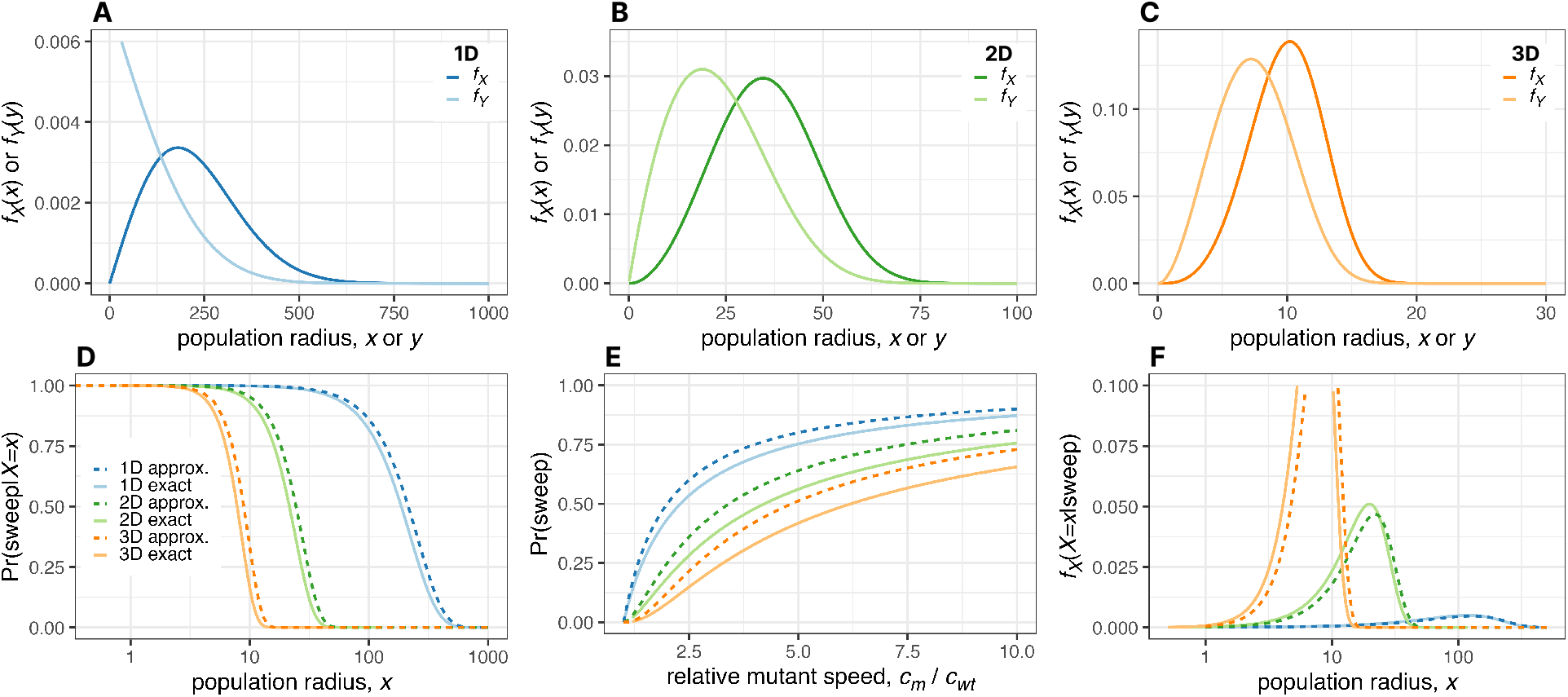
Analytical and numerical solutions of the macroscopic model. **A.-C**. Probability density of the wildtype population radius at the time the first surviving mutant arises (dark curves) and of the distance between the origins of the mutant and wildtype expansions (light curves) in one dimension **(A)**, two dimensions **(B)** and three dimensions **(C). D**. Approximate and exact solutions for the sweep probability conditioned on the wildtype population radius. **E**. Approximate and exact solutions for the unconditional sweep probability. **F**. Approximate and exact solutions for the probability density of the wildtype population radius at the time the first surviving mutant arises, conditioned on this mutant achieving a selective sweep. Except where parameter values are explicitly varied, we set *c*_wt_ = 0.15,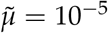, *ρ* = 0.23 and *c*_m_ = 0.31. Formulas for all these curves are summarised in Table S2.

#### Location of the first mutant

Next, we compute the probability density for the distance *Y* between the first surviving mutant and the centre of the wildtype population. Since mutants arise in proportion to the number of wildtype cells, we have *f*_*Y*_(*y*|*X* = *x*) *dy ∝ D*(*y*), where *D*(*y*) is the number of cells at distance *y. D*(*y*) corresponds to the infinitesimal shell, *D*(*y*) = 4*πy*^2^ *dy* **1***{y ≤ x}*, where the last term is an indicator function that defines the boundary of the wildtype population. After normalisation, we obtain

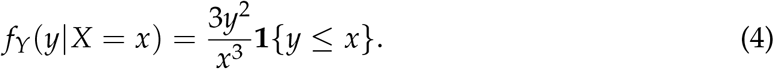

We calculate the unconditional probability density of *Y* by marginalizing out *X*,

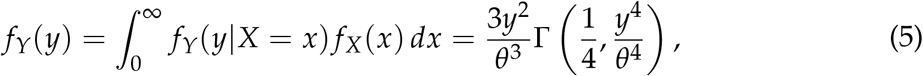

where 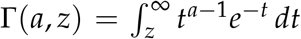 is the incomplete gamma function. We then find E[*Y*] *≈* 0.68 *θ* and Var[*Y*] *≈* 0.070 *θ*^2^. Similar results pertain in one and two dimensions (SI text; Figure 2A-C).

#### Conditional sweep probability

To compute the sweep probability, we first need an expression for the remaining wildtype population, *N*_wt_. Therefore, we introduce time measure *τ*, with *τ* = 0 when the first mutant arises. Recall that *t* = 0 at the origin of the wildtype population, so *t* = *τ* + *x*/*c*_wt_. Once the mutant population starts expanding, we have 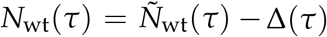, where *Δ*(*τ*) is the number of wildtype cells that the mutant has replaced, and 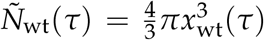 is the wildtype population size had there been no mutant. While the mutant is surrounded by the wildtype, we have 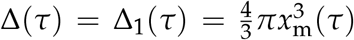. After the mutant breaches the wildtype boundary, *Δ*(*τ*) is given by the intersection formula of two balls, which we denote *Δ*_2_(*τ*) (see SI text for the explicit formula). Together, we have

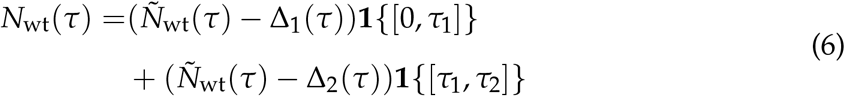

where *τ*_1_ is the time at which the mutant reaches the wildtype boundary and *τ*_2_ is the time at which the mutant has entirely replaced the wildtype. Noting that

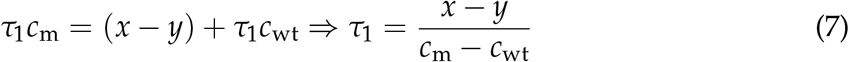

and

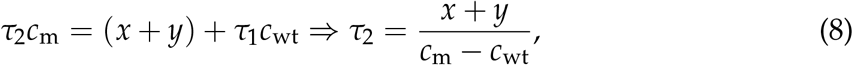

we express *N*_wt_(*τ*) in terms of *x* and *y* (see SI text). We then apply Claim 1 to obtain the conditional sweep probability,

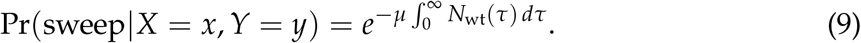

Although this integral can be solved analytically (SI text), the resulting expression is complicated and not very enlightening, and we therefore seek simpler approximations. An especially fruitful approach is to assume *y* = 0, so that *f*_*Y*_(*y*|*X* = *x*) = *δ*(*y*), where *δ* is the Dirac delta function. This yields an upper bound on the conditional sweep probability because, due to geometrical symmetry, the time required for a mutant to sweep (*τ*_2_ in eqn. 8) is minimal when *y* = 0. With this approximation, eqn. 9 simplifies drastically as 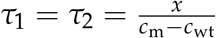, and we do not need to integrate over *Δ*_2_(*τ*). The analytic solution is

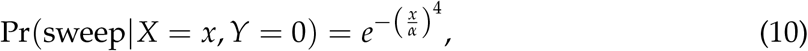

where 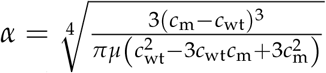 is a characteristic length. Despite being based on a seemingly crude approximation, eqn. 10 is remarkably close to the exact conditional sweep probability (Figure 2D). The corresponding approximations in one and two dimensions (SI text) are likewise simple and useful.

#### Unconditional sweep probability

We now have all the necessary ingredients to compute the unconditional sweep probability by marginalizing out *X* and *Y* from the conditional sweep probability,

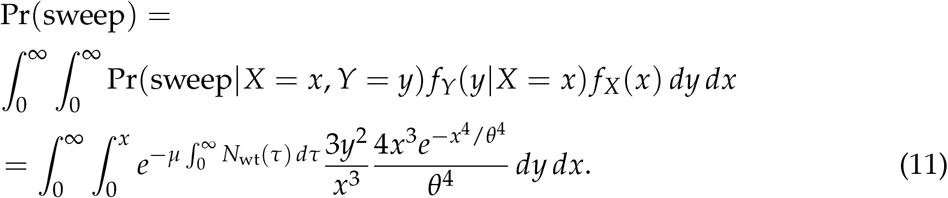

As before, we use the approximation *y* = 0 to obtain the strikingly simple result

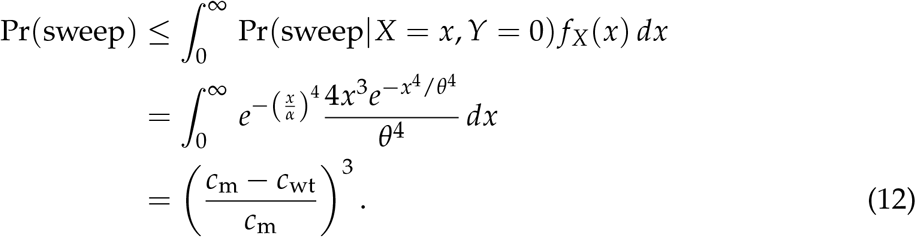

This result generalizes to analogous one- and two-dimensional models, with the exponent 3 replaced by the respective spatial dimension (SI text). The approximate expressions are close to the exact solution in one dimension, and to numerical evaluations of the integral in two and three dimensions (Figure 2E). The sweep probability is independent of the mutation rate not only when *y* = 0 but also in the general case of eqn. 11. This follows from a more general result (SI text).

When the expanding wildtype must displace a resident competitor (as in our agent-based simulations, to follow), the sweep probability can be approximated in terms of evolutionary parameters via the FKPP solution as

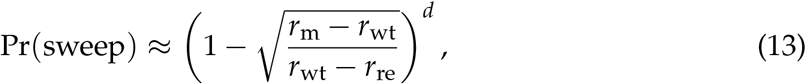

where *r*_m_, *r*_wt_ and *r*_re_ are the proliferation rates of the mutant invader, wildtype invader and resident populations, respectively, and *d* is the spatial dimension.

#### Conditional arrival time of the first mutant

In biological systems where it is infeasible to track evolutionary dynamics, selective sweeps must be inferred from subsequent genetic data. For example, we might observe a public mutation in a tumour and ask when this mutation occurred. We can use our model to obtain the probability distribution of the radius *X* at the time the first mutant arose, given that we observe a selective sweep, simply by applying Bayes’ theorem,

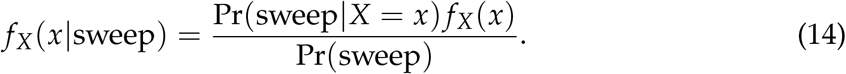

Using the approximation *y* = 0, we obtain the Weibull distribution

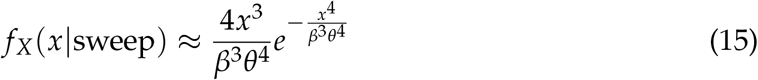

where 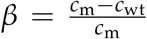. This approximation and the corresponding results in one and two dimensions are close to the exact solutions (Figure 2F).

### Simulations

To gauge the robustness of our theoretical predictions against stochastic growth and discretization, we measured the frequency of selective sweeps in a two-dimensional agent-based model.

#### An agent-based realisation of the two-dimensional spatial Moran process

We suppose that the wildtype population invades a habitat initially occupied by a resident competitor, which is a plausible biological scenario for both invasive species [40] and invasive tumours [28, 32]. Our agent-based model thus has three types of individuals: resident, wildtype invader, and mutant invader with local proliferation rates *r*_re_, *r*_wt_ and *r*_m_. Localised competition between wildype and resident slows the wildtype expansion and creates potential for selective sweeps. We implemented this model using the *demon* agent-based modelling framework [41] within the *warlock* computational workflow [42], which facilitates running large numbers of simulations on a high-performance computing cluster. We have previously applied the same framework to studying cancer evolution [28, 42]. Further model details are given in Materials and Methods.

The agent-based simulations provide useful insights additional to the macroscopic model because, although the general setup is the same, they differ in several ways. Space in the simulations is divided into discrete patches; the times between birth and dispersal events are exponentially distributed random variables constituting another source of stochasticity; population boundaries are rough, not smooth; the expansion wave front is typically not sharp, and changes shape as the wave progresses; and the mutant will have increased propagation speed when competing with the resident population. Hence we would not expect perfect agreement between the results of the two models.

#### Linking the microscopic and macroscopic models

Our agent-based model approximately resembles a spatial death-birth Moran process (also known as the stepping stone model) [43, 44]. Expansion speeds in the spatial Moran process can in turn be approximated using the Fisher-Kolmogorov-Petrovsky- Piscounov (FKPP) equation [37, 38], which predicts that the mutant will expand within the wildtype with constant radial expansion speed *c*_m_ dependent on the difference in their local proliferation rates *a*_m_ = *r*_m_ *− r*_wt_ [33, 34, 36]. Analogously, we have a constant expansion speed of the wildtype *c*_wt_ dependent on *a*_wt_ = *r*_wt_ *− r*_re_. To compare the results of our discrete-space simulations to our continuous-space macroscopic model, we measured the propagation speeds of the wildtype within the resident, and of the mutant within the wildtype (see Materials and Methods). Further investigations of propagation speeds in this model will be the subject of a further study.

#### Simulation results

Given the considerable differences between the models, the probability density functions resulting from the macroscopic and microscopic models are reassuringly consistent. The radius at the time the first surviving mutant arises is slightly lower in the simulations than in our analytical model (Figure 3A). Similarly, the distribution for the location of the first surviving mutant coincides well except for a small offset of the mean (Figure 3B). Such offsets are expected due to discretization effects and the fact that, in the simulations, the propagation front needs to be established before the expansion can proceed. The sweep probabilities in simulations are nevertheless very close to our analytical predictions and change very little when varying the mutation rate over orders of magnitude, confirming our prediction that the sweep probability is independent of the mutation rate (Figure 3C; Figures S1-S4).

**Figure 3.**
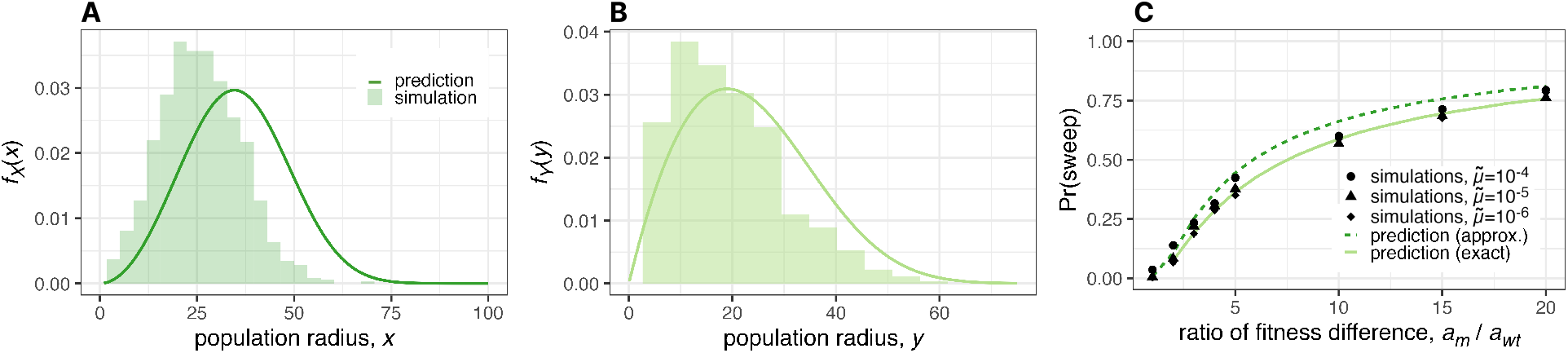
Agent-based simulation results versus macroscopic model predictions. **A**. Probability density of the wildtype population radius at the time the first surviving mutant arises. **B**. Probability density of the distance between the wildtype origin and the location at which the first successful mutant is born. **C**. Unconditional selective sweep probability versus the ratio of the proliferation rate differences *a*_wt_ = *r*_n_ *− r*_re_ and *a*_m_ = *r*_m_ *− r*_wt_. Each histogram in panels **A** and **B** and each data point in **C** is based on 1,000 replicates. Except where parameter values are explicitly varied, we set *m* = 0.05,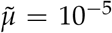, *K* = 16, *r*_re_ = 0.91, *r*_wt_ = 1 and *r*_m_ = 1.3, leading to speeds *c*_wt_ *≈* 0.15 and *c*_m_ *≈* 0.31 and mutant survival probability *ρ ≈* 0.23. Results for alternative parameter values and less stringent sweep definitions are shown in Figures S1-S3. The conversion from simulation parameters to macroscopic model parameters is explained in Materials and Methods.

### Comparison with other growth models

To further demonstrate the robustness of our findings, we compare them to results obtained from alternative models. Instead of permitting mutants to grow within the wildtype population, one might instead assume that dispersal is restricted to the wildtype population boundary. In this case, a complete sweep is impossible and we instead ask whether the first arising mutant envelopes the wildtype. Interestingly, this envelopment probability is independent of the mutation rate and obtains values close to the sweep probabilities of our main model [21] (Figures 4 and S5; SI text). Alternatively, one might consider exponential growth with no competition, and ask whether a single mutant comes to dominate the population. Adapting our methods to this case (see SI text), we find that a single mutant is unlikely to become dominant unless its exponential growth rate is several times larger than that of the wildtype (Figure S6). Although our focus is on expanding populations, we also applied our methods to compute the sweep probability in constant populations (SI text), obtaining a result that depends on both mutation rate and population size (Figure 4), in agreement with prior analyses [45, 46].

**Figure 4.**
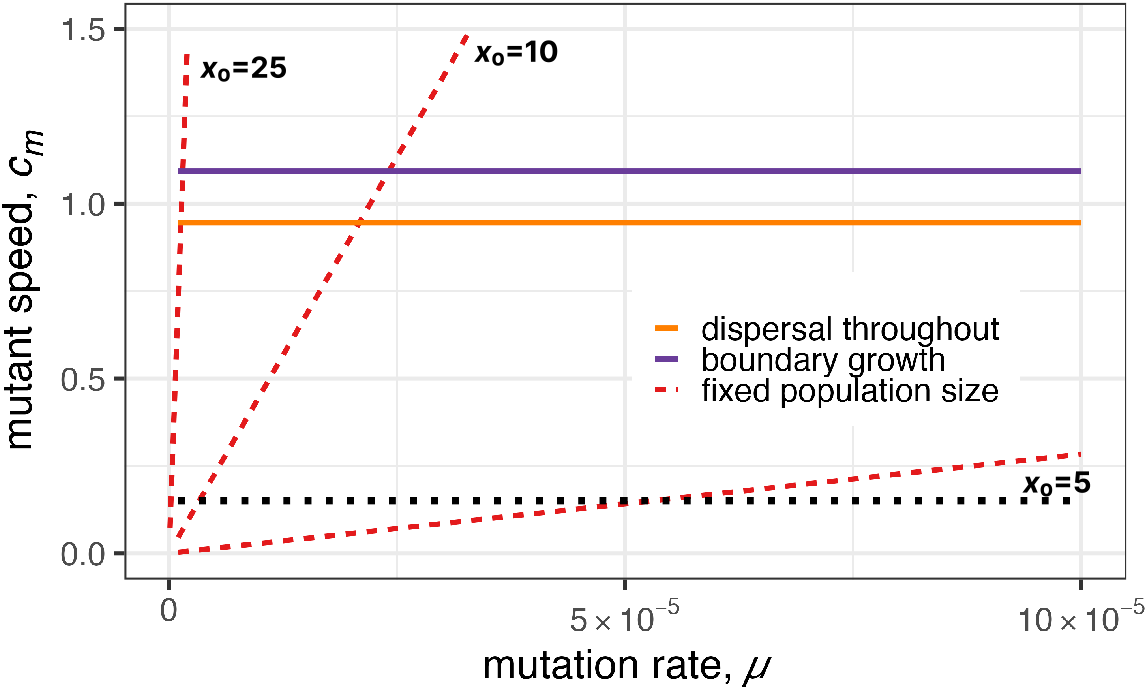
Comparison of sweep probabilities between different growth models in 3D. We show the tuples (*μ, c*_m_) that yield Pr(sweep) = 0.5 for models with proliferation restricted to the boundary (solid purple line), population size fixed at radius *x*_0_ (dashed red lines), and our main model in which mutants can expand within the expanding wildtype population (solid orange line). In the two range expansion models, the wildtype speed is set to *c*_wt_ = 0.15 (dotted black line).

### Application to cancer

Our findings have especially interesting implications for understanding cancer evolution. Here we consider a wildtype tumour that evolves while invading a resident population of normal cells.

#### Arrival of the first driver mutation during tumour growth

Our model has only three parameters: the mutation rate conditional on survival, *μ*, and the wildtype and mutant propagation speeds, *c*_wt_ and *c*_m_. To estimate *c*_wt_ for human tumours, we follow a similar procedure to prior studies [21, 47]. Consider a tumor that grows to volume *V* between 1 and 10 cm^3^ in time *T* between 5 and 20 years. The propagation speed can then be estimated as

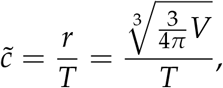

which equates to between 1 and 40 *μ*m per day. Given that the diameter of a typical cancer cell is *l ≈* 20 *μ*m [48] and the generation time (cell cycle time) is *τ*_*G*_ *≈* 4 days [49], we can switch units to obtain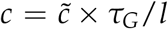, which is between 0.15 and 7 cell diameters per generation.

The rate of acquiring advantageous (driver) mutations is usually estimated at around 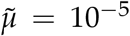 per cell per generation [47, 50]. For the survival probability *ρ*, we assume values between 0.09 and 0.5, in agreement with inferred values in colorectal tumours [51].

Together, these parameter values lead to a typical length scale

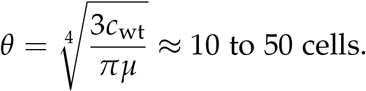

The expected values of *X* and *Y* are then E[*X*] = 9 to 45 cell diameters and E[*Y*] = 7 to 35 cell diameters. It follows that a tumour, having acquired sufficient mutations to grow, will likely gain further driver mutations already during early development.

#### Ubiquity of clonal interference in cancer

During tumour growth, we predict very low sweep probabilities unless the mutant grows much faster within the tumour than the tumour grows into surrounding tissue (Figure 2E, 3C). When sweeps do occur, they will begin and end in the very early stages of tumour growth. Assuming *c*_m_ is at most ten times *c*_wt_, we find that the expected tumour radius when a selective sweep occurred, given that a sweep did occur, is no more than 50 cells, corresponding to a population size *N* of no more than 400,000 cells. The time for the sweep to be completed is then *τ*_2_ *≈* 40 generations, at a population size of approximately 800, 000 cells. These numerical estimates are highly insensitive to varying parameter values because 𝔼[*X*], 𝔼[*Y*] and the expected value of *f*_*X*_(*x*|sweep) are proportional to the characteristic length *θ*, which varies with the fourth root of *μ* and *c*_wt_. Hence, in three dimensions, changing *μ* or *c*_wt_ by a factor of 100 changes our estimates by only a factor of 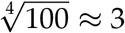.

## DISCUSSION

Here we have used mathematical modelling and analysis to determine the expected frequency of classic selective sweeps versus incomplete or soft sweeps during range expansions. We find that this frequency is generally expected to be low, even for mutations with very high selection coefficients. Moreover, when the wildtype and mutant radial expansion speeds are constant, the sweep probability can be expressed solely in terms of those speeds, which can in turn be related, through the FKPP equation or other standard models, to the selection coefficient, dispersal rates, and other basic parameters. Why is the sweep probability independent of the mutation rate? An intuitive explanation is that if the mutation rate is higher then, on the one hand, the first advantageous mutation is likely to arise in a smaller population – meaning that it has less distance to travel to achieve a sweep – but, on the other hand, competing mutations will also tend to arise sooner than in the case of a lower mutation rate. These two effects exactly cancel out under the assumption of constant radial growth speeds.

We make three arguments to justify our focus on this particular model of range expansions. First, our model corresponds to the continuum approximation of standard mathematical models of range expansions and spatial population genetics: the spatial Moran process (or stepping stone model [44]) and the biassed voter model (which is equivalent to an Eden growth model extended to allow local dispersal and competition throughout the population) [36]. These models are well understood, intuitive, and easy to parameterise. Second, because our model is relatively permissive to selective sweeps, it provides useful upper bounds for selective sweep probabilities in more complex scenarios, as further explained below.

Our third justification is that, in much the same way as the Moran and Wright-Fisher processes are the most useful, tractable models of evolution in constant-sized, non-spatial populations, so the constant-radial-speed model yields the clearest results for range expansions. Haldane’s famous rule of thumb is that the fixation probability of a weakly beneficial allele in a large, non-spatial population of constant size is approximately proportional to its relative advantage in terms of proliferation rate [52]. Here we have obtained the comparably simple result that the probability of a strongly beneficial allele achieving a selective sweep in an expanding population is approximately equal to its relative advantage in terms of radial expansion speed, raised to the power of the spatial dimension. For instance, if the radial expansion speed at which a mutant spreads within the wildtype population is twice the speed at which the wildtype expands then the probability of this mutant achieving a selective sweep can be approximated simply as (1 *−* 1/2)^2^ = 1/4 in two dimensions and 1/8 in three dimensions.

Some alternative models have been considered previously. Antal and colleagues [21] used a macroscopic model similar to ours to investigate the case in which mutations arise only at the boundary of a range expansion. Given that selective sweeps are then impossible, the interesting outcome is when the mutant envelops the wildtype. Ralph & Coop [45] and Martens and colleagues [20, 46] instead considered constant-sized populations, and found that selective sweeps are likely only if the population width is much smaller than a characteristic length scale, which depends on the mutation rate, the dispersal rate, the effective local population density, and the strength of selection. In SI Text we compare our results to those of these prior studies and consider other alternative growth laws. The general conclusions are that selective sweeps are predicted to be rare except in small populations and that our model provides a useful upper bound on the sweep probability in range expansions. Selective sweeps will be even less frequent when the wildtype radial expansion speed increases over time (as is typical in the middle stage of tumour progression) or when mutant expansion is restricted to the tumour boundary (which may be the case in some tumours [28, 29, 53]).

A selective sweep can occur only if the rate of spread of an advantageous mutation exceeds the expansion speed of the wildtype population. This scenario is plausible, for example, in a biofilm growing slowly under antibiotic stress, in which case our findings predict the evolutionary dynamics of antibiotic resistance. Our results apply equally to an invasive species that is still adapting to its new conditions and whose range expansion is slowed by the need to modify its environment (niche construction) or by interspecific interactions [40]. If the invader must outcompete a resident then the sweep probability can be approximated via the FKPP solution in terms of proliferation rates as in eqn. 13. The dimensional exponent in this result implies that selective sweeps are most likely to occur in species invading essentially linear habitats, such as coastlines.

Our three-dimensional results are particularly relevant to understanding the nature of solid tumour evolution. There are two ways in which tumours might acquire the multiple clonal drivers (advantageous mutations) observed in sequencing studies. The first route is via sequential population bottlenecks. Suppose that early tumour development comprises not one but several successive range expansions. Growth repeatedly stalls due to constraints such as hypoxia, immune control, and physical barriers. Driver mutations enable subclones to escape these constraints and invade new territory, each time purging genetic diversity so that the final, prolonged expansion originates from a single highly transformed cell. This episodic model is conventional and has been particularly well characterised in recent studies of colorectal cancer [54, 55] and breast cancer [56]. The second way to acquire clonal drivers is via selective sweeps during the final range expansion. Our results suggest that the first process is the more important. Selective sweeps of even extremely strong drivers are highly unlikely to occur in tumours containing more than a few hundred thousand cells, equivalent to less than a cubic millimeter in volume. Pervasive genetic heterogeneity and parallel evolution [57] are therefore revealed to be straightforward consequences of spatial expansion, expected to be ubiquitous across solid tumours, irrespective of mutation rate and degree of genomic instability.

## METHODS

### Numerical Integration

We performed numerical integration using the MATLAB function ‘trapz’. For values of *x* close to 0 and beyond 2*θ, f*_*X*_(*x*) is relatively very small and its contribution to the integrals is negligible. Hence, to avoid numerical errors without compromising precision, we set the lower and upper bounds of integration to 0.001*θ* and 3*θ* when integrating over *f*_*X*_(*x*). Similarly, we set 0.001*θ* as the lower bound of the integration over *y*. We set interval widths to 0.001*θ* for *x ≤* 0.5, and 0.01*θ* for *x* > 0.5.

### Agent-based simulations

Individuals in our agent-based model are subdivided into well-mixed demes on a regular two-dimensional grid. The demes have identical carrying capacities, *K*, and are initially filled with residents, except that a single wildtype invader is introduced at the centre of the grid. At each time step, an individual is chosen at random, with probability weighted by fitness, to be replaced by two offspring. Each offspring then either migrates, with probability *m*, to a neighbouring deme in a randomly chosen direction, or remains in its parent deme. Local density dependence is implemented by imposing a very high death rate whenever a deme is above carrying capacity. Mutation is coupled to wildtype reproduction. To improve computational efficiency, resident individuals are not permitted to disperse; we verified with additional simulations (to be published in a later study) that this asymmetry has only a minor effect on the wildtype expansion speed. Further model details have been published previously [28, 42]. All simulations were performed on City, University of London’s Hyperion cluster.

For all simulations, we set the deme size to *K* = 16 and the migration probability *m* = 0.05. For such small *m*, the probability to survive drift can be estimated as the probability to become locally fixed in one deme. Since the within-deme dynamics approximate a Moran process, this probability is 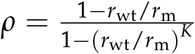, where *r*_wt_ and *r*_m_ are the proliferatio rates [58]. We measured propagation speeds by simulating the expansion of a wildtypeinto a large resident population, or of a mutant into a large wildtype population, in the absence of mutation and then applying linear regression to the growth curve of the effective radius, defined as the square root of the population size divided by *π*. For each parameter set, we took the mean speed from ten replicates, for which the standard deviation was consistently below 2% of the mean. We set the proliferation rates to *r*_re_ = 0.91 and *r*_wt_ = 1.0 for the resident and wildtype, respectvively, leading to *a*_wt_ = *r*_wt_ *− r*_re_ = 0.09. The measured wildtype speed was then *c*_wt_ *≈* 0.15. We varied the mutant proliferation rate *r*_m_ from 1.1 to 3.0, so that *a*_m_ = *r*_m_ *− r*_wt_ ranged from 0.1 to 2.0. The measured mutant speeds are presented in Table I.

**Table 1.** Correspondence between difference in proliferation rates *a*_m_ and mutant propagation speed *c*_m_, measured in simulations.

|  |  |  |  |  |  |  |  |
| --- | --- | --- | --- | --- | --- | --- | --- |
| $a_{\text{m}}$ | 0.1 | 0.2 | 0.3 | 0.4 | 0.5 | 0.6 | 0.7 |
| $c_{\text{m}}$ | 0.14 | 0.23 | 0.31 | 0.38 | 0.46 | 0.53 | 0.60 |
| $a_{\text{m}}$ | 0.8 | 0.9 | 1.0 | 1.1 | 1.2 | 1.3 | 1.4 |
| $c_{\text{m}}$ | 0.68 | 0.75 | 0.82 | 0.89 | 0.96 | 1.03 | 1.11 |
| $a_{\text{m}}$ | 1.5 | 1.6 | 1.7 | 1.8 | 1.9 | 2.0 | |
| $c_{\text{m}}$ | 1.17 | 1.24 | 1.31 | 1.39 | 1.46 | 1.52 | |

## AUTHOR CONTRIBUTIONS

RN conceived the research question and supervised the project. RN and AS designed the research. AS and RK carried out the mathematical analysis. MB performed agent-based simulations. All authors wrote and approved the manuscript.

## ACKNOWLEDGMENTS

AS was supported by the European Union’s Horizon 2020 research and innovation programme under the Marie Składowska-Curie EvoGamesPlus grant agreement no. 955708. MB was supported by an award from the City, University of London Research Pump-priming Fund. RN was supported by the National Cancer Institute of the National Institutes of Health under Award Number U54CA217376. The content is solely the responsibility of the authors and does not necessarily represent the official views of the National Institutes of Health.

## CODE AVAILABILITY

The code and data needed to reproduce our results can be found at https://zenodo.org/records/10775693.

## Notes

https://zenodo.org/records/10775693

### Competing Interest Statement

The authors have declared no competing interest.

### Summary of Updates

Reverted the title. Revised the abstract and first paragraph of the Introduction to emphasize how our work differs from prior research. Revised the second paragraph of the Introduction to summariz ongoing debate about the mode of tumour evolution. Split the second paragraph of the Discussion into two paragraphs and added a couple of sentences comparing our rule of thumb to Haldane's.

