## Supplementary material for "Selective sweep probabilities in spatially expanding populations": Supplemenary text, figures and tables

### Supporting Information Text

#### 1. Proof of Claim 1

*Proof of claim 1.* By definition of the per capita mutation rate, we have $\mu N(t_1)dt$ successful mutations in the infinitesimal time interval $[t_1, t_1 + dt]$. We sum over the time interval $[0, t]$ and obtain $\lambda = \int_0^t \mu N(t)\, dt$. We assume that mutations occur independently of each other (infinite alleles) such that the stochastic process is a non-homogeneous Poisson process with mean $\lambda$. □

#### 2. Connection between per-capita and per-division mutation rate

In our macroscopic model, we assume a constant per-capita mutation rate $\tilde{\mu}$. Often, it is assumed that mutations are coupled to division. Here, we show how a per-capita mutation rate connects to a per-division mutation rate. Suppose that after each division one of the two daughter cells obtains a driver mutation with probability $p_\mu$. (The probability that both daughter cells obtain a driver mutation scales with $p_\mu^2$, which can be neglected for sufficiently small $p_\mu$.) Suppose cells proliferate at rate $b$ and die at rate $d$. Those rates may depend on the time or location. If division rates are constant over time and space, then the per-division mutation probability $p_\mu$ yields a constant per-capita mutation rate $\tilde{\mu} = p_\mu b$. The death rate may still differ in time and space.

#### 3. Exact solution for the conditional sweep probability

Here, we provide the full expression and exact solution for the conditional sweep probability $\Pr(\text{sweep}|X = x, Y = y)$ for the three-dimensional case (eqn. 9 in the main text). From the main text, we have

$$\Pr \text{sweep}|X = x, Y = y = e^{-\mu \int_0^t N_{\text{wt}}(\tau)\, d\tau}. \qquad [1]$$

were $N_{\text{wt}}$ is the size of the remaining wildtype population and following eqn. 6 from the main text the integral can be divided into

$$\int_0^\infty N_{\text{wt}}(\tau)\, d\tau = \int_0^{\tau_1} \tilde{N}_{\text{wt}}(\tau) - \Delta_1(\tau)\, d\tau + \int_{\tau_1}^{\tau_2} \tilde{N}_{\text{wt}}(\tau) - \Delta_2(\tau)\, d\tau. \qquad [2]$$

Here $\tilde{N}_{\text{wt}}$ and $\Delta_1(m)$ are the population sizes of the wildtype and mutant in case of undisturbed radial growth such that

$$\begin{aligned} \tilde{N}_{\text{wt}} &= \frac{4}{3}\pi x_{\text{wt}}^3(\tau) \\ \Delta_1(\tau) &= \frac{4}{3}\pi x_{\text{m}}^3(\tau). \end{aligned} \qquad [3]$$

Furthermore, $\Delta_2(\tau)$ is given by the intersecting volume of two balls with radii $x_{\text{wt}}$ and $x_{\text{m}}$ at distance $y$. Following ref. (1), we have

$$\Delta_2(\tau) = \frac{\pi}{12y}(x_{\text{wt}} + x_{\text{m}} + y)^2(y^2 + 2yx_{\text{m}} - 3x_{\text{m}}^2 + 2yx_{\text{wt}} + 6x_{\text{wt}}x_{\text{m}} - 3x_{\text{wt}}^2). \qquad [4]$$

We can express $x_{\text{wt}}$ and $x_{\text{m}}$ in terms of the wildtype radius at the time the first mutant occurred $x$ and the time $\tau$ from the emergence of the first mutant using that

$$\begin{aligned} x_{\text{wt}}(\tau) &= c_{\text{wt}}t = c_{\text{wt}}(\tau + x/c_{\text{wt}}) \\ x_{\text{m}}(\tau) &= c_{\text{m}}\tau. \end{aligned} \qquad [5]$$

Now, we are ready to integrate over $N_{\text{wt}}(\tau)$ and obtain

$$\int_0^\infty N_{\text{wt}}(\tau)\, d\tau = \frac{\pi\left(15x^4\left(3c_{\text{m}}^2 - 3c_{\text{m}}c_{\text{wt}} + c_{\text{wt}}^2\right) - y^4\left(23c_{\text{m}}^2 + 31c_{\text{m}}c_{\text{wt}} + 23c_{\text{wt}}^2\right) - 210c_{\text{m}}^2x^2y^2\right)}{45(c_{\text{m}} - c_{\text{wt}})^3}. \qquad [6]$$

Inserting this result into eqn. 1 gives us the conditional sweep probability. The exact solution reduces to the solution in the main text with $y = 0$, for which

$$\int_0^\infty N_{\text{wt}}(\tau)\, d\tau = \frac{1}{\mu}\left(\frac{x}{\alpha}\right)^4 \quad \text{with} \quad \alpha = \sqrt[4]{\frac{3(c_{\text{m}} - c_{\text{wt}})^3}{\pi\mu(c_{\text{wt}}^2 - 3c_{\text{wt}}c_{\text{m}} + 3c_{\text{m}}^2)}}. \qquad [7]$$

#### 4. The sweep probability is independent of the mutation rate

We show that the unconditional selective sweep probability $\Pr(\text{sweep})$ (eqn. 11 in the main text) is independent of the mutation rate. We start by defining homogeneous functions and stating a lemma for homogeneous functions. We then show how this lemma proves independence of the unconditional sweep probability and eventually prove the lemma. Here, we focus on the three-dimensional case whereas the independence for the two- and one-dimensional case is shown later.

#### A. Homogeneous functions.

**Definition 1.** *A function $f : \mathbb{R}^n \to \mathbb{R}$ is called homogeneous of degree $k$ if*

$$f(sx_1, sx_2, ..., sx_n) = s^k f(x_1, x_2, ..., x_n) \quad [8]$$

*for every $x_1, x_2, ..., x_n$ and $s \neq 0$.*

Noteworthy, polynomials in which all terms have the same polynomial degree $k$ are homogeneous functions with degree $k$. For example, $5x^2y^2 + \pi x^4$ is homogeneous in $x$ and $y$ with degree 4.

**Lemma 1.** *Let $\nu > 0$. Let $H(x, y)$, $P(x, y)$ and $Q(x, y)$ be homogeneous functions in $x$ and $y$ with degrees $h, p, q > 0$ respectively and $Q(x, y) \neq 0$ for all $x > 0$ and $y > 0$. Then, the integral*

$$I = \int_0^\infty \int_0^x e^{-\nu H(x,y)} \frac{P(x, y)}{Q(x, y)} \, dy \, dx \quad [9]$$

*is proportional to $\nu^{\frac{q-p-2}{h}}$.*

**B. The sweep probability is independent of the mutation rate.** To apply the lemma on the unconditional sweep probability, we set $\nu = \mu$ and

$$\begin{aligned} H(x, y) &= \int_0^\infty N_{\text{wt}}(\tau) \, d\tau + \frac{3c_{\text{wt}}}{\pi} x^4 \\ P(x, y) &= 3y^2 \times 4x^3 \\ Q(x, y) &= \frac{3c_{\text{wt}}}{\pi} x^3. \end{aligned} \quad [10]$$

It is straightforward that $P(x, y)$ and $Q(x, y)$ are homogeneous in $x$ and $y$ with degrees $p = 5$ and $q = 3$. Looking at eqn. 6, it is also apparent that $H(x, y)$ is homogeneous with degree 4. Using those expressions and applying lemma 1, we have

$$\Pr(\text{sweep}) = \mu \times I \propto \mu \times \mu^{\frac{3-5-2}{4}} = \mu^0, \quad [11]$$

which proves the independence of the mutation rate.

We found that the approximate solution using $f_Y(y|X = x) = \delta(y)$ is also independent of the mutation rate indicating a more general result. Looking at the derivation above, this becomes clear. Consider fixed equations for $\Pr(sweep|X = x, Y = y)$ and $f_X(x)$. For any density function $f_Y(y|X = x)$ that is homogeneous in $x$ and $y$ with degree -1, we can define functions $P(x, y)$ and $Q(x, y)$ such that their difference in degrees is $p - q = -2$. After application of the lemma, it follows independence.

#### C. Proof of Lemma 1.

*Proof of lemma 1.* We show the relation by making the substitutions

$$\hat{x} = \nu^{1/h} x \quad \text{and} \quad \hat{y} = \nu^{1/h} y$$

such that

$$\begin{aligned} H(\hat{x}, \hat{y}) &= \nu H(x, y) \\ P(\hat{x}, \hat{y}) &= \nu^{p/h} P(x, y) \\ Q(\hat{x}, \hat{y}) &= \nu^{q/h} Q(x, y) \end{aligned}$$

and

$$dx = \nu^{-1/h} d\hat{x} \quad \text{and} \quad dy = \nu^{-1/h} d\hat{y}.$$

Inserting the substitutions in eqn. 9, we obtain

$$\begin{aligned} I &= \int_0^\infty \int_0^x e^{-\nu H(x,y)} \frac{P(x, y)}{Q(x, y)} \, dy \, dx \\ &= \int_0^\infty \int_0^{\hat{x}} e^{-H(\hat{x},\hat{y})} \frac{\nu^{-p/h} P(\hat{x}, \hat{y})}{\nu^{-q/h} Q(\hat{x}, \hat{y})} \nu^{-1/h} \nu^{-1/h} \, d\hat{y} \, d\hat{x} \\ &= \nu^{-2/h} \nu^{(q-p)/h} \int_0^\infty \int_0^{\hat{x}} e^{-H(\hat{x},\hat{y})} \frac{P(\hat{x}, \hat{y})}{Q(\hat{x}, \hat{y})} \, d\hat{y} \, d\hat{x} \end{aligned}$$

The remaining integral is no longer dependent on parameter $\nu$ and shows the desired scaling with $\nu^{\frac{q-p-2}{h}}$. □

#### 5. Sweep probability in one dimension

We consider a population that is expanding in one dimension in two directions with constant speed. If $x_{\mathrm{wt}}$ describes the length from the origin of the population to the leading edge, then the population size is $N_{\mathrm{wt}} = 2x_{\mathrm{wt}}(t) = 2c_{\mathrm{wt}}t$. We follow the derivations described in the main text.

**A. Arrival time and location of the first mutant.** We start by computing the probability density for the time at which the first mutant arises. Using claim 1, we compute the probability that no mutant occurs until time $t$ to be

$$P_0 = e^{-\mu \int_0^t 2c_{\mathrm{wt}} t' dt'} = e^{-\mu c_{\mathrm{wt}} t^2} = e^{-\left(\frac{t}{\kappa_{1\mathrm{D}}}\right)^2} \quad \text{with} \quad \kappa_{1\mathrm{D}} = \sqrt{\frac{1}{\mu c_{\mathrm{wt}}}}. \tag{12}$$

The probability density function for the timing of the first surviving mutant is then

$$f_T(t) = \frac{d(1-P_0)}{dt} = \frac{2t}{\kappa_{1\mathrm{D}}^2} e^{-\left(\frac{t}{\kappa_{1\mathrm{D}}}\right)^2} \tag{13}$$

With a change of variables $t = \frac{x}{c_{\mathrm{wt}}}$, we obtain the probability density for the population radius $X$ at the time the first surviving mutant arises, which is

$$f_X(x) = \frac{2x}{\theta_{1\mathrm{D}}^2} e^{-\left(\frac{x}{\theta_{1\mathrm{D}}}\right)^2} \quad \text{with} \quad \theta_{1\mathrm{D}} = \sqrt{\frac{c_{\mathrm{wt}}}{\mu}}. \tag{14}$$

Since all distances $y$ from the wildtype origin have the same amount of proliferating cells, the probability density for the location of the first surviving mutant $Y$ conditional on the wildtype radius $X = x$ is a homogeneous. After consideration of boundary conditions and normalization, we have

$$f_Y(y|X=x) = \frac{1}{x} 1\{y \le x\}, \tag{15}$$

The unconditional probability density of $y$ is computed by

$$f_Y(y) = \int_0^\infty f_X(x) f_Y(y|X=x)\, dx = \frac{1}{\theta_{1\mathrm{D}}} \Gamma\left(\frac{1}{2}, \frac{y^2}{\theta_{1\mathrm{D}}^2}\right). \tag{16}$$

**B. Exact sweep probability.** In one dimension, we obtain a closed-form expression for the exact solution of the sweep probability. Suppose a mutant is born at distance $y$ once the wildtype radius reached length $x$. Again, we set $\tau = 0$ at the time the mutant is born. Following the main text, the remaining wildtype population is given by

$$N_{\mathrm{wt}} = (2x_{\mathrm{wt}} - 2x_{\mathrm{m}})\, \mathbf{1}[0, \tau_1] + (x_{\mathrm{wt}} - x_{\mathrm{m}} + y)\, \mathbf{1}[\tau_1, \tau_2] \tag{17}$$

where $\tau_1 = \frac{x-y}{c_{\mathrm{m}} - c_{\mathrm{wt}}}$ is the time at which the mutant population reaches the first leading edge of the wildtype and $\tau_2 = \frac{x+y}{c_{\mathrm{m}} - c_{\mathrm{wt}}}$ is the time at which the mutant reaches the second leading edge of the wildtype such that the sweep is completed. Next, we substitute $x_{\mathrm{wt}} = x + c_{\mathrm{wt}}\tau$ and $x_{\mathrm{m}} = c_{\mathrm{m}}\tau$ and then apply claim 1 to compute the probability that no further mutant arises before the sweep is completed, which leads to

$$\Pr(\text{sweep}|X=x, Y=y) = e^{-\mu \int_0^\infty N_{\mathrm{wt}}(\tau)\, d\tau} = e^{-\frac{x^2+y^2}{\alpha_{1\mathrm{D}}^2}} \quad \text{with} \quad \alpha_{1\mathrm{D}} = \sqrt{\frac{c_{\mathrm{m}} - c_{\mathrm{wt}}}{\mu}}. \tag{18}$$

Then, we marginalize out $Y$, which gives us

$$\Pr(\text{sweep}|X=x) = \int_0^\infty \Pr(\text{sweep}|X=x, Y=y) f_Y(y|X=x)\, dy = \frac{\sqrt{\pi}\alpha_{1\mathrm{D}}}{2x} e^{-\left(\frac{x}{\alpha_{1\mathrm{D}}}\right)^2} \operatorname{erf}\left(\frac{x}{\alpha_{1\mathrm{D}}}\right), \tag{19}$$

where $\operatorname{erf}(x) = \frac{2}{\sqrt{\pi}} \int_0^x e^{-t^2}\, dt$ is the Gaussian error function. The exact formula for the unconditional sweep probability is obtained by marginalizing out $X$. The result is

$$\Pr(\text{sweep}) = \int_0^\infty \Pr(\text{sweep}|X=x) f_X(x)\, dx = \frac{\beta' \cot^{-1}\left(\sqrt{1+\beta'}\right)}{\sqrt{1+\beta'}}, \tag{20}$$

where $\beta' = \frac{c_{\mathrm{m}} - c_{\mathrm{wt}}}{c_{\mathrm{wt}}}$ is the speed difference relative to the wildtype speed. We notice that the exact solution for the sweep probability is independent of the mutation rate.

**C. Approximate sweep probability.** We use the approximation explained in the main text and set $y = 0$, which translates into a probability density $f_Y(y|X = x) = \delta(y)$. Under this assumption, the conditional sweep probability is given by

$$\Pr(\text{sweep}|X = x) = \Pr(\text{sweep}|X = x, Y = 0) = e^{-\left(\frac{x}{\alpha_{1\mathrm{D}}}\right)^2}. \quad [21]$$

and the unconditional sweep probability becomes

$$\Pr(\text{sweep}) = \int_0^\infty \Pr(\text{sweep}|X = x) f_X(x)\, dx = \frac{c_\mathrm{m} - c_\mathrm{wt}}{c_\mathrm{m}}. \quad [22]$$

Using Bayes theorem, we compute the probability distribution of $X$ given a sweep has occurred,

$$f_X(X = x|\text{sweep}) = \frac{2x}{\theta_{1\mathrm{D}}^2 \beta} e^{-\frac{x^2}{\theta_{1\mathrm{D}}^2 \beta}}, \quad [23]$$

where $\beta = \frac{c_\mathrm{m} - c_\mathrm{wt}}{c_\mathrm{m}}$. The distribution of $X$ is still described by a Weibull distribution with shape parameter 2. The difference is only a scaling factor of the characteristic length $\theta_{1\mathrm{D}} \to \theta_{1\mathrm{D}}\sqrt{\beta}$.

#### 6. Sweep probability in two dimensions

We consider a population that is expanding radially in two dimensions at constant speed. If $x_\mathrm{wt}$ is the radius of the wildtype population, then the population size is $N = \pi x_\mathrm{wt}^2 = \pi(c_\mathrm{wt} t)^2$. We follow the derivations in the main text.

**A. Arrival time and location of the first mutant.** We start and compute the probability density for the time at which the first mutant arises. Using claim 1, we compute the probability that no mutant occurs until time $t$ to be

$$P_0 = e^{-\mu \int_0^t \pi c_\mathrm{wt}^2 t'^2\, dt'} = e^{-\mu \frac{\pi}{3} c_\mathrm{wt}^2 t^3} = e^{-\left(\frac{t}{\kappa_{2\mathrm{D}}}\right)^3} \quad \text{with} \quad \kappa_{2\mathrm{D}} = \sqrt[3]{\frac{3}{\mu \pi c_\mathrm{wt}^2}}. \quad [24]$$

Taking the derivative gives us the probability density for the arrival time of the first surviving mutant,

$$f_T(t) = \frac{d(1 - P_0)}{dt} = \frac{3t^2}{\kappa_{2\mathrm{D}}^3} e^{-\left(\frac{t}{\kappa_{2\mathrm{D}}}\right)^3}. \quad [25]$$

We perform a change of variables $t = \frac{x}{c_\mathrm{wt}}$ to obtain the probability density for the radius of the wildtype population $X$ at the time the first mutant arises that is

$$f_X(x) = \frac{3x^2}{\theta_{2\mathrm{D}}^3} e^{-\left(\frac{x}{\theta_{2\mathrm{D}}}\right)^3}, \quad \text{with} \quad \theta_{2\mathrm{D}} = \sqrt[3]{\frac{3c_\mathrm{wt}}{\pi\mu}}. \quad [26]$$

To compute the probability density for the distance $Y$ between the first mutant and the centre of the wildtype population, we notice that there at distance $y$, there are $2\pi y\, dy$ dividing cells at distance $y$. Given that the first mutant occurred when the wildtype population had radius $X = x$, we have $f_Y(y|X = x) \propto y^2$. After taking care of the boundary condition and normalization, we get

$$f_Y(y|X = x) = \frac{2y}{x^2} 1\{y \leq x\}, \quad [27]$$

The unconditional probability density of $y$ is obtained by

$$f_Y(y) = \int_0^\infty f_Y(y|X = x) f_X(x)\, dx = \frac{2y}{\theta_{2D}^2} \Gamma\left(\frac{1}{3}, \frac{y^3}{\theta_{2D}^3}\right). \quad [28]$$

**B. Exact sweep probability.** We start by finding the expression for the remaining wildtype population $N_\mathrm{wt}$ given that $X = x$ and $Y = y$. As in the three- and one-dimensional cases, we have the remaining wildtype population given by two formulas such that

$$N_\mathrm{wt}(\tau) = (\tilde{N}_\mathrm{wt}(\tau) - \Delta_1(\tau))\mathbf{1}\{[0, \tau_1]\} + (\tilde{N}_\mathrm{wt}(\tau) - \Delta_2(\tau))\mathbf{1}\{[\tau_1, \tau_2]\} \quad [29]$$

with

$$\begin{aligned} \tilde{N}_\mathrm{wt}(\tau) &= \pi x_\mathrm{wt}^2 \\ \Delta_1(\tau) &= \pi x_\mathrm{m}^2 \end{aligned} \quad [30]$$

given by the area of discs with radii $x_\mathrm{wt}$ and $x_\mathrm{m}$ and

$$\begin{aligned} \Delta_2(\tau) =& x_\mathrm{wt}^2 \cos^{-1}\left(\frac{y^2 + x_\mathrm{wt}^2 - x_\mathrm{m}^2}{2yx_\mathrm{wt}}\right) + x_\mathrm{m}^2 \cos^{-1}\left(\frac{y^2 + x_\mathrm{m}^2 - x_\mathrm{wt}^2}{2yx_\mathrm{m}}\right) \\ &- \frac{1}{2}\Big((-y + x_\mathrm{wt} + x_\mathrm{m})(y + x_\mathrm{wt} - x_\mathrm{m})(y - x_\mathrm{wt} + x_\mathrm{m})(y + x_\mathrm{wt} + x_\mathrm{m})\Big)^{1/2}. \end{aligned} \quad [31]$$

is the intersecting area of two circles with radii $x_{\mathrm{wt}}$, $x_{\mathrm{m}}$ at distance $y$ that we took from ref. (2). The formulas for the times $\tau_1$ and $\tau_2$ remain the same to the one- and two-dimensional case that are $\tau_1 = \frac{x-y}{c_{\mathrm{m}}-c_{\mathrm{wt}}}$ and $\tau_2 = \frac{x+y}{c_{\mathrm{m}}-c_{\mathrm{wt}}}$. Again, we replace $x_{\mathrm{wt}} = x + c_{\mathrm{wt}}\tau$ and $x_{\mathrm{m}} = c_{\mathrm{m}}\tau$.

Applying claim 1 on the remaining wildtype population, we obtain the conditional sweep probability

$$\Pr(\mathrm{sweep}|X = x, Y = y) = \int_0^\infty \Pr(\mathrm{sweep}|X = x, Y = y), \quad [32]$$

which can be solved numerically. The unconditional sweep probability is then given by the integral

$$\begin{aligned}\Pr(\mathrm{sweep}) &= \int_0^\infty \int_0^\infty \Pr(\mathrm{sweep}|X = x, Y = y) f_Y(y|X = x) f_X(x)\, dy\, dx,\\ &= \int_0^\infty \int_0^x e^{-\mu \int_0^\infty N_{\mathrm{wt}}(\tau)d\tau} \frac{2y}{x^2} \frac{3x^2 e^{-x^2/\theta_{2\mathrm{D}}^3}}{\theta_{2\mathrm{D}}^3}\, dy\, dx,\end{aligned} \quad [33]$$

which we solved numerically.

**C. Independence of the mutation rate.** We apply lemma 1 on the integral form of Pr(sweep) by defining $\nu = \mu$ and

$$\begin{aligned}H(x,y) &= \int_0^\infty N_{\mathrm{wt}}(\tau)d\tau + x^3 \frac{3c_{\mathrm{wt}}}{\pi},\\ P(x,y) &= 2y \times 3x^2,\\ Q(x,y) &= x^2 \frac{\pi}{3\mathrm{cwt}},\end{aligned} \quad [34]$$

which are homogeneous and have degrees $h = 3$, $p = 3$ and $q = 2$. It follows that

$$\begin{aligned}\Pr(\mathrm{sweep}) &= \mu \times \int_0^\infty \int_0^x e^{-\mu H(x,y)} \frac{P(x,y)}{Q(x,y)}\, dy\, dx\\ &\propto \mu \times \mu^{\frac{2-3-2}{3}} = \mu^0,\end{aligned} \quad [35]$$

which proves independence of he mutation rate.

**D. Approximate sweep probability.** We use the approximation explained in the main text and set $y = 0$, which translates into a probability density $f_Y(y|X = x) = \delta(y)$. Under this assumption, the conditional sweep probability is given by

$$\Pr(\mathrm{sweep}|X = x) = \Pr(\mathrm{sweep}|X = x, Y = 0) = e^{-\left(\frac{x}{\alpha_{2\mathrm{D}}}\right)^3}, \quad \text{with} \quad \alpha_{2\mathrm{D}} = \sqrt[3]{\frac{3(c_{\mathrm{m}} - c_{\mathrm{wt}})^2}{\pi\mu(2c_{\mathrm{m}} - c_{\mathrm{wt}})}} \quad [36]$$

and the unconditional sweep probability becomes

$$\Pr(\mathrm{sweep}) = \left(\frac{c_{\mathrm{m}} - c_{\mathrm{wt}}}{c_{\mathrm{m}}}\right)^2 \quad [37]$$

Using Bayes theorem, we compute the probability distribution of $X$ given a sweep has occurred,

$$f_X(X = x|\mathrm{sweep}) = \frac{3x^2}{\theta_{2\mathrm{D}}^3 \beta^2} e^{-\frac{x^3}{\theta_{2\mathrm{D}}^3 \beta^2}} \quad [38]$$

where $\beta = \frac{c_{\mathrm{m}} - c_{\mathrm{wt}}}{c_{\mathrm{m}}}$. The distribution of $X$ is still described by a Weibull distribution with shape parameter 3. The difference is only a scaling factor of the characteristic length $\theta_{2\mathrm{D}} \to \theta_{2\mathrm{D}}\beta^{\frac{2}{3}}$.

#### 7. Sweep probabilities in other growth models

We focused on constant radial expansion speeds for multiple reasons explained in the main text. However, to obtain more generality we also computed sweep probabilities in other growth models. Here, we consider (i) constant populations in which the mutant propagates with constant speed (ii) radial growth of the wildtype and the mutant where proliferation is restricted the population boundary and (iii) exponential growth. We briefly discuss (iv) sigmoidal growth leading to Lotka-Volterra type of dynamics.

**A. Constant populations.** We consider a constant population of spherical form with radius $x_0$ leading to total population size $N_0 = \frac{4}{3}\pi x_0^3$. The probability distribution of the wildtype radius $X$ at the time the fist mutant arises can be formally written as the delta function,

$$f_X(x) = \delta(x - x_0). \quad [39]$$

The probability density for the distance $Y$ between the first mutant and the centre of the wildtype population conditioned on $X = x$ is the same as for expanding populations. However, we can replace the random variable $X = x$ by the fixed radius $x_0$ such that

$$f(y) = f(y|X = x_0) = \frac{3y^2}{x_0^3}\mathbf{1}\{y < x_0\}. \quad [40]$$

Next, we calculate the sweep probability conditioned on $X$ and $Y$. We denote the remaining wildtype population once a mutant arose by $N_{\mathrm{wt}}(\tau)$ and analogously to the previous calculation introduce the time measure $\tau$ that starts with the emergence of the first mutant. We have

$$N_{\mathrm{wt}} = (\tilde{N}_{\mathrm{wt}}(\tau) - \Delta_1(\tau))\mathbf{1}\{[0, \tau_1]\} + (\tilde{N}_{\mathrm{wt}}(\tau) - \Delta_2(\tau))\mathbf{1}\{[\tau_1, \tau_2]\} \quad [41]$$

where the terms are identical to those described in section 3 except for constant wildtype radius $x_0$ such that

$$\begin{aligned} \tilde{N}_{\mathrm{wt}} &= \frac{4}{3}\pi x_0^3 \\ \Delta_1(\tau) &= \frac{4}{3}\pi x_{\mathrm{m}}^3 \\ \Delta_2(\tau) &= \frac{\pi}{12y}(x_0 + x_{\mathrm{m}} + y)^2(y^2 + 2yx_{\mathrm{m}} - 3x_{\mathrm{m}}^2 + 2yx_0 + 6x_0x_{\mathrm{m}} - 3x_0^2) \end{aligned} \quad [42]$$

where $x_{\mathrm{m}} = x_{\mathrm{m}}(\tau) = c_{\mathrm{m}}\tau$. Using claim 1, the conditional sweep probability is computed by

$$\Pr(\text{sweep}|Y = y) = e^{-\mu\int_0^\infty N_{\mathrm{wt}}(\tau)\,d\tau} \quad \text{with} \quad \int_0^\infty N_{\mathrm{wt}}(\tau)\,d\tau = \frac{\pi\left(45x_0^4 - 210x_0^2y^2 - 23y^4\right)}{45c_{\mathrm{m}}}. \quad [43]$$

Integrating the conditional sweep probability together with $f_Y(y)$ does not yield a closed-form expression.

For analytical insight, let us assume again that the mutant originates in the centre of the wildtype population, $f_Y(y) = \delta(y)$. We then obtain

$$\Pr(\text{sweep}) = \Pr(\text{sweep}|Y = 0) = e^{-\frac{\mu\pi x_0^4}{c_m}}. \quad [44]$$

Because the timing of the first mutation plays no role in constant populations, our reasoning for the independence of the mutation rate no longer holds, and we obtain a sweep probability dependent on the mutation rate. We compared this sweep probability for different $x_0$ to sweep probabilities in expanding populations (Figure 4 and S5).

Performing the same analysis in one and two dimensions, we obtain

$$\begin{aligned} \Pr(\text{sweep}) &= e^{-\frac{\mu x_0^2}{2c_{\mathrm{m}}}} \quad \text{in 1D} \\ \Pr(\text{sweep}) &= e^{-\frac{2\mu\pi x_0^3}{3c_{\mathrm{m}}}} \quad \text{in 2D} \end{aligned} \quad [45]$$

**Comparison with Ralph & Coop:** Motivated by parallel adaptation on the species level, Ralph & Coop investigated selective sweeps in constant populations using a similar approach (3). However, whereas we assume spherical wildtype populations, Ralph & Coop consider a general shapes and use scaling arguments. To derive a concrete expression, they ignore boundary effects and obtain “the expected number of other mutations to arise in an area of diameter $a$ in the time it takes the wave to cover that area". Simplifying eqn. (4) in ref. (3) and keeping their notation, we have

$$\frac{2a^3\lambda}{v} \quad \text{in 1D} \quad \text{and} \quad \frac{\pi a^3\lambda}{v} \quad \text{in 2D}. \quad [46]$$

The number of mutations arising in an area over time is a Poisson process (claim 1). Thus, we can interpret this number as mean of a Poisson distribution. A selective sweep has occurred if no other mutation arose and thus the sweep probability reads

$$\begin{aligned} \Pr(\text{sweep}) &= P_0 = e^{-\frac{2a^2\lambda}{v}} \quad \text{in 1D}, \\ \Pr(\text{sweep}) &= P_0 = e^{-\frac{\pi a^3\lambda}{v}} \quad \text{in 2D}. \end{aligned} \quad [47]$$

We identify $v$ as the speed of the mutant $c_{\mathrm{m}}$, $\lambda$ as the local mutation rate conditioned on survival $\mu = \tilde{\mu}\rho$ and $a$ as the maximal travelled distance by the first mutant, which is $x_0$ in the case we set $y = 0$. Finally, we find the solution of Ralph & Coop (eqn. 47) to be identical to our solution (eqn. 45) up to multiplication by a constant that can be explained by differing boundary conditions. The conclusion in either case is that full selective sweeps are highly unlikely for sufficiently large population radius $x_0$.

**Comparison with Martens et al.:** Martens and colleagues investigated the likelihood of clonal interference and its impact on the speed of evolution in spatially structured constant-size populations (4), and then studied the implications for understanding cancer initiation (5). Two modes of evolution are considered: (i) acquisition via subsequent selective sweeps and (ii) acquisition with parallel arising mutations which interfere with each other. To distinguish between these two modes, the authors compare the timescale for a surviving mutation to occur, $t_{\mathrm{mut}}$, and the timescale for a mutation to sweep through the entire constant wildtype population, $t_{\mathrm{fix}}$. Equating these two timescales leads to a critical length of the wildtype population:

$$\begin{aligned} L_c &= \left(\frac{c_0}{2s_0\mu}\right)^{\frac{1}{2}} \quad \text{in 1D} \\ L_c &= \left(\frac{c_0}{2s_0\mu}\right)^{\frac{1}{3}} \quad \text{in 2D.} \end{aligned} \tag{48}$$

The authors argue that clonal interference is very likely in populations of size $L \gg L_c$, whereas we should expect selective sweeps when $L \ll L_c$. To compare this result with our model, we put $L_c$ into eqn. 45. Therefore, we match parameters by $c_0 \leftrightarrow c_{\mathrm{m}}$, $2s\mu \leftrightarrow \rho\tilde{\mu} = \mu$ and $x_0 \leftrightarrow L$ leading to

$$\begin{aligned} \Pr(\text{sweep}) &= e^{-\frac{x_0^2}{2L_c^2}} \quad \text{in 1D} \\ \Pr(\text{sweep}) &= e^{-\frac{\pi x_0^3}{3L_c^3}} \quad \text{in 2D.} \end{aligned} \tag{49}$$

Indeed, we have $\Pr(\text{sweep}) \to 0$ for $x_0 \gg L_c$ and $\Pr(\text{sweep}) \to 1$ for $x_0 \ll L_c$. We conclude that the result of Martens et al. are in agreement with our result for constant population sizes.

**B. Exponential growth.** We consider an exponentially growing population, $N(t) = e^{rt}$. Applying claim 1, the probability that no mutants occur until time $t$ is given by

$$P_0 = e^{-\mu \int_0^t e^{rt'}\,dt'} = e^{-\frac{\mu}{r}(e^{rt}-1)}. \tag{50}$$

Dropping the $-1$, this equation coincides with the solution for the stochastic birth-death process obtained by Durrett (eqn. [25] in Ref. (6)) up to the constant $V_0$ that is caused by fluctuations in early growth history and is neglected within our deterministic growth assumption. We proceed to calculate the probability density of $T$ by

$$f(t) = \frac{d(1-P_0}{dt} = \mu e^{rt} e^{-\frac{\mu}{r}e^{rt}}, \tag{51}$$

Instead of asking for the radius, it is more sensible to ask for the population size of the wildtype at the arrival of the first mutant. We write $N_x = e^{rt}$, such that $t = \frac{1}{r}\ln(N_x)$. After substitution, we have

$$f_{N_x}(N_x) = \frac{\mu}{r} e^{-\frac{\mu}{r}N_x}, \tag{52}$$

which is an exponential distribution. The probability for the radius $X$ can then be calculated by assuming spherical growth starting from one cell, $N_x = \frac{4}{3}\pi x + 1$. For mathematical convenience, we neglect the $+1$ term. The probability distribution of $X$ reads

$$f_X(x) = \frac{3x^2}{\theta_{\exp}^3} e^{-\frac{x^3}{\theta_{\exp}^3}} \quad \text{with} \quad \theta_{\exp} = \sqrt[3]{\frac{3r}{4\pi\mu}}. \tag{53}$$

Noteworthy, exponential growth does not come with an underlying spatial model and migration of the mutant is unclear. Nevertheless, no such assumptions are required to compute the location at which the first mutant arises. The conditional probability distribution of $Y$ remains the same to our main model that is

$$f_Y(y|X=x) = \frac{3y^2}{x^3} 1\{y \le x\}. \tag{54}$$

To obtain the unconditional probability for $y$, we can marginalize out $X$ giving us

$$\begin{aligned} f_Y(y) &= \int_0^\infty f_Y(y|X=x) f_X(x)\,dx \\ &= \frac{9y^2}{\theta_{\exp}^3} \int_y^\infty \frac{e^{-\frac{x^3}{\theta_{\exp}^3}}}{x}\,dx \\ &= \frac{3y^2}{\theta_{\exp}^3} \mathrm{Ei}\left(\frac{x^3}{\theta_{\exp}^3}\right), \end{aligned} \tag{55}$$

where $\mathrm{Ei}(z) = \int_0^\infty \frac{1}{t} e^{-t}\,dt$ is te exponential integral function.

In the exponential growth model, there is no competition such that the wildtype population will never go extinct and the mutant will never become fixed. We can however compute whether the mutant reaches a certain frequency $\chi$ before another mutant arises. We assume deterministic growth of the wildtype and the mutant and random time for the emergence of the mutant. We have

$$\chi = \frac{N_\mathrm{m}}{N_\mathrm{m} + N_\mathrm{wt}}. \qquad [56]$$

Now, define $t_1$ to be the time at which the first mutant occurs and let $t_2$ be the time at which the mutant has grown to frequency $\chi$. Then

$$\chi = \frac{e^{r_\mathrm{m} t_2}}{e^{r_\mathrm{m} t_2} + e^{r_\mathrm{wt}(t_1+t_2)}}. \qquad [57]$$

Here, $t_1$ is a random variable that follows the probability density described by eqn. 51, and $t_2$ can be computed by solving the equation for $\chi$ giving us

$$t_2 = \frac{r_\mathrm{wt} t_1 - \ln\left(\frac{1-\chi}{\chi}\right)}{r_\mathrm{m} - r_\mathrm{wt}}. \qquad [58]$$

The conditional sweep probability (or more precisely the probability for the first mutant to reach frequency $\chi$ without interference of another mutant) is given by

$$\Pr(\text{sweep}|T = t_1) = e^{-\lambda} \quad \text{with} \quad \lambda = \mu \int_0^{t_2} e^{r_\mathrm{wt}(t_1+\tau)}\, d\tau = \frac{\mu}{r_\mathrm{wt}} e^{r t_1} \left(e^{r_\mathrm{wt} t_2} - 1\right), \qquad [59]$$

where $t_2 = t_2(t_1)$ is a function of $t_1$ described above. Eventually, the unconditional sweep probability is computed by

$$\begin{aligned} \Pr(\text{sweep}) &= \int_0^\infty \Pr(\text{sweep}|T = t_1) \times f_T(t_1)\, dt_1 \\ &= \int_0^\infty e^{-\frac{\mu}{r_\mathrm{wt}} e^{r t_1}\left(e^{r_\mathrm{wt} t_2(t_1)} - 1\right)} \times \mu e^{r t_1} e^{-\frac{\mu}{r} e^{r t_1}}\, dt_1 \end{aligned} \qquad [60]$$

which we solved numerically and present in Figure S6. Notably, the probability is dependent on the mutation rate.

**C. Boundary growth without proliferation in the interior.** Antal and colleagues studied aspects of the evolutionary dynamics in boundary-driven growth where cell proliferation is restricted to the boundary only (7). More properties of this system have further been studied extensively (8). Whereas we assume turnover of the entire population, Antal et al. consider turnover only at the boundary and focus on the three-dimensional case. A full selective sweep is impossible in this model since individuals located away from the boundary neither proliferate nor die. Instead, we can consider the probability that a mutant will envelop the wildtype and thus becomes the only proliferating population. Using this interpretation, eqn. (17) in ref. (7) provides the unconditional sweep probability that is

$$\Pr \text{sweep} = \frac{9 + \beta^2}{\beta^2} \frac{2}{1 + e^{3\pi/\beta}}, \qquad [61]$$

with $\beta = \sqrt{v^2 - 1}$ and we can identify $v = \frac{c_\mathrm{m}}{c_\mathrm{wt}}$. The unconditional sweep probability is independent of the mutation rate just like in our model. Furthermore, Antal et al. find similar expressions for the arrival time of the first mutant $f_T(t)$ (eqn. 15 in (7)) and the conditional sweep probability $\Pr(\text{sweep}|X = x)$ (eqn. 16 in ref. (7)) as well as the size of the wildtype population when the first mutant occurs (eqn. 25 in ref. (7) provides the cumulative density function). The unconditional sweep probability in the case of boundary restricted proliferation is lower compared to the sweep probability allowing proliferation throughout the tumor including the interior (Figure 4 and S5).

**D. Sigmoidal growth.** Many biological systems do not grow infinitely but reach a carrying capacity. There are a number of common models described by differential equations that start with initially experiential growth that eventually converges to a carrying capacity (e.g. logistic, Gompertzian (9)). The most common sigmoidal growth model is the logistic growth. If there are at least two types of populations with different growth parameters, the logistic growth model is naturally extended to a competitive Lotka-Volterra system. Those models have long been investigated and closed-form expressions are usually not known. Furthermore, the implementation of competition can take multiple forms. A fitness advantage in Lotka-Volterra systems can be obtained by an increased growth rate, an increased carrying capacity or frequency dependent competition factors that are subject of studies in evolutionary game theory (e.g. (10)). A study of selective sweeps in sigmoidal growth models thus comes with further complications that cannot be solved with methods presented in this paper alone.

Nevertheless, until the first mutant arises, there is only one population type (and thus no complicated competition between different types). Thus, we can compute the time and location of the first surviving mutant. We consider a population that grows logistically according to $N(t) = \frac{K}{1+Ke^{-rt}}$. We apply claim 1, and obtain the probability no mutation occurs until time $t$,

$$P_0 = e^{-\frac{\mu K}{r} \ln\left(\frac{e^{rt}+K}{K+1}\right)}. \qquad [62]$$

The probability density is then obtained taking the derivative,

$$f_T(t) = \frac{d(1-P_0)}{dt} = \frac{\mu K(K+1)^{\frac{\mu K}{r}} e^{rt}}{(e^{rt}+K)^{\frac{\mu K}{r}+1}}. \quad [63]$$

The probability density for the population size $N_x$ at the time the first mutant arises can be obtained by substituting $N_x = \frac{K}{1+Ke^{-rt}}$ yielding

$$f_{N_x}(N_x) = \frac{\mu\,(K+1)^{\frac{\mu K}{r}}\,(K-N_x)^{\frac{\mu K}{r}-1}}{rK^{\frac{2\mu K}{r}-1}}. \quad [64]$$

Assuming spherical growth, $N_x = \frac{4}{3}\pi x^3$, the probability density for the radius $X$ can be obtained by substituting $N_x = \frac{4}{3}\pi x^3$. We obtain

$$f_X(x) = \frac{\mu\,(K+1)^{\frac{\mu K}{r}}\left(K - \frac{4\pi x^3}{3}\right)^{\frac{\mu K}{r}-1} 4\pi x^2}{rK^{\frac{2\mu K}{r}-1}}. \quad [65]$$

The distance $Y$ of the first mutant from the wildtype's origin conditioned on the radius $X = x$ remains the same as before,

$$f_Y(y|X=x) = \frac{3y^2}{x^3}1\{y \le x\}. \quad [66]$$

To obtain the unconditional density for $Y$, we can marginalize out $X$,

$$f_Y(y) = \int_0^\infty f_Y(y|X=x)f_X(x)\,dx. \quad [67]$$

which gives us

$$\frac{12\pi y^2\mu(K+1)^{\frac{\mu K}{r}}}{rK^{\frac{2\mu K}{r}-1}} \int_y^\infty \frac{\left(K - \frac{4\pi x^3}{3}\right)^{\frac{\mu K}{r}-1}}{x}\,dx. \quad [68]$$

The integral can be evaluated numerically.

**E. Sweep probability at 50% .** To compare the models in the space of parameters $(\mu, c_\mathrm{m})$, we can compute the tuples $(\mu, c_\mathrm{m})$ at which the sweep probability is 50%. The tuples can be expressed as curve $c_\mathrm{m}(\mu)$ for which $\Pr(\text{sweep}) = 0.5$ leading to the following expressions for the three-dimensional expansions:

- Growth throughout (eqn.[14] in the main text): $c_\mathrm{m}(\mu) \approx 6.3\, c_\mathrm{wt}$
- Boundary proliferation (eqn. 61): $c_\mathrm{m}(\mu) \approx 7.2\, c_\mathrm{wt}$
- constant population (eqn. 44): $c_\mathrm{m}(\mu) = \frac{\mu\pi x_0^4}{\ln(2)}$.

We solved the first two expressions numerically. The plots are shown in the main text (Figure 4).

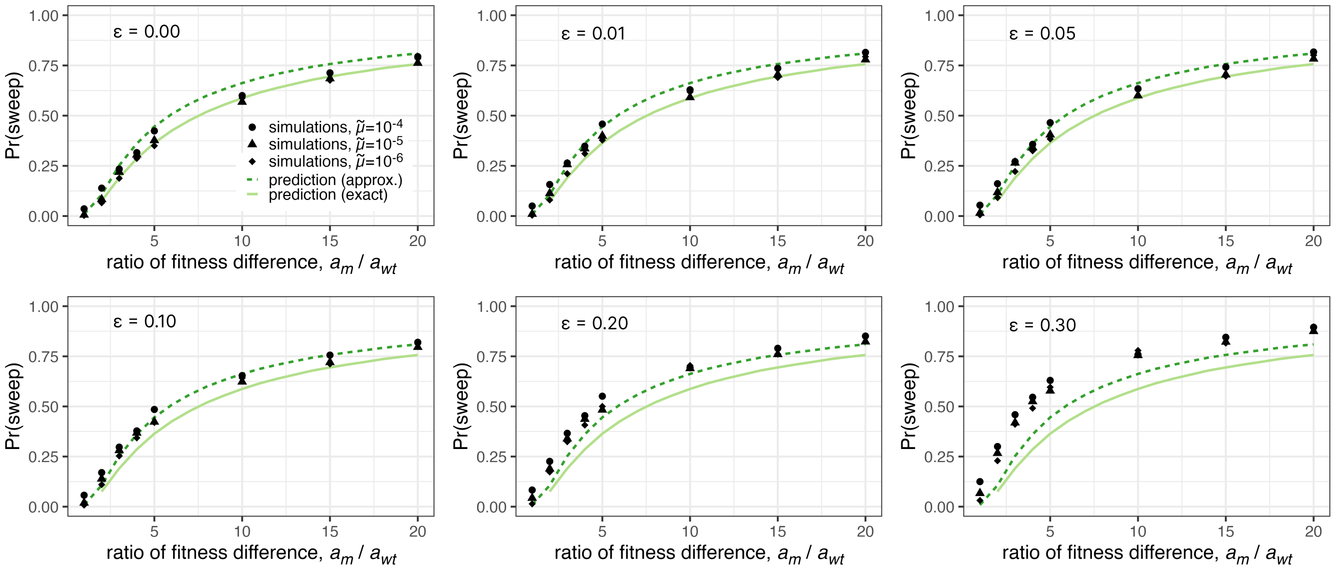


**Fig. S1.** Simulation results of selective sweep frequencies with different cutoffs in 2D. A selective sweep might be incomplete because of other minority populations. Here, we show simulation results using a weakened definition of a selective sweep. A selective sweep is said to be successful if the mutant's frequency reaches $1-\epsilon$. The case $\epsilon = 0.00$ corresponds to a complete selective sweep as shown and discussed in the main text. Parameters were set to $m = 0.05$, $K = 16$, $\tilde{\mu} = 10^{-4}, 10^{-5}, 10^{-6}$, $r_{\text{re}} = 0.91$, $r_{\text{wt}} = 1.0$. We vary over $r_{\text{m}}$ between $1.1$ and $3.0$ corresponding to different $a_{\text{m}}$. Conversion between simulation and macroscopic model parameters is described in the Materials and Methods.

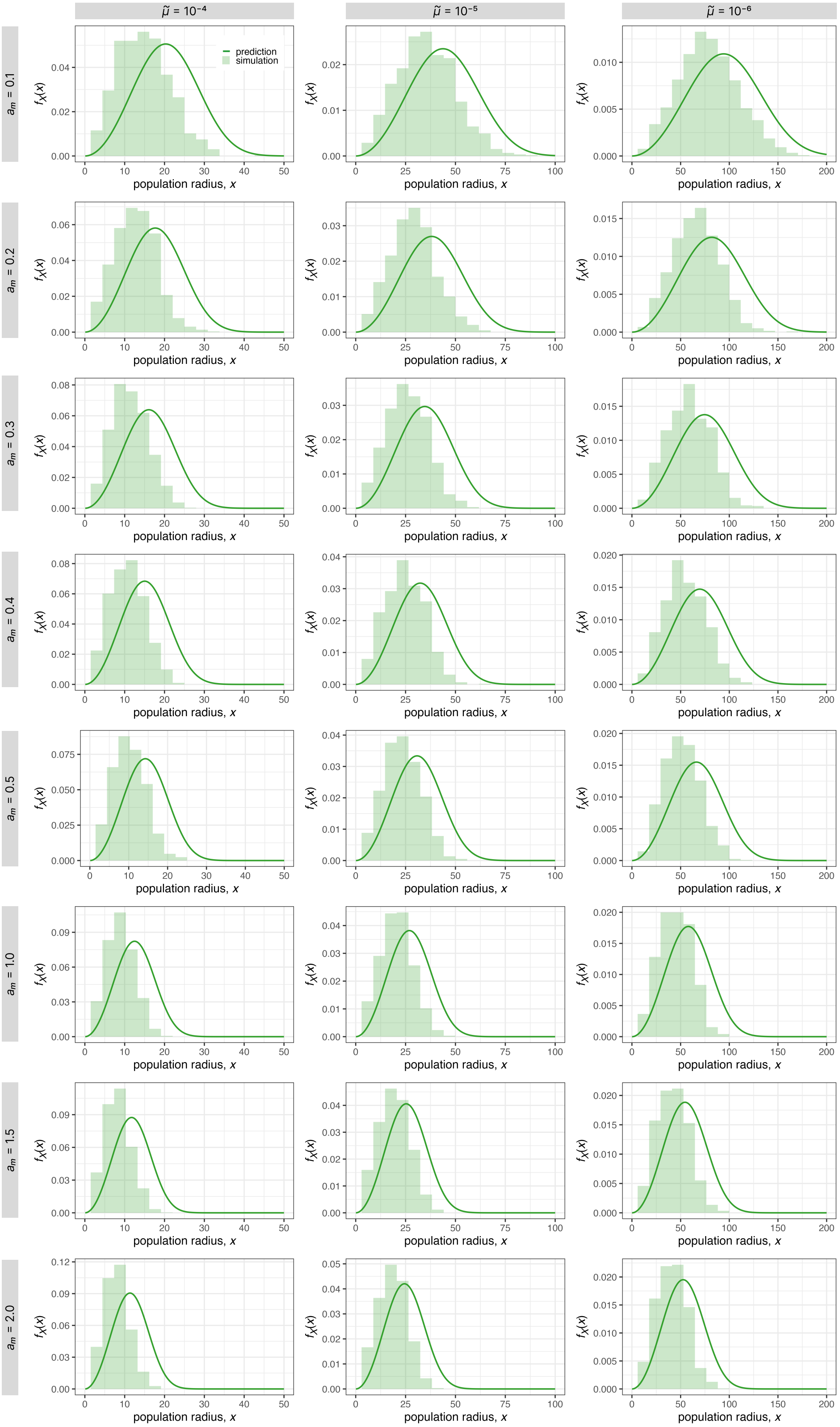


**Fig. S2.** Comparison of simulation results with our analytic predictions of $f_X(x)$ in 2D for different parameter combinations. We use $m = 0.05$, $K = 16$, $r_{\text{re}} = 0.91$, $r_{\text{wt}} = 1.00$. We vary over the mutation rate $\mu$ and $r_{\text{m}}$ corresponding to different $a_m$. Conversion between simulation and macroscopic model parameters is described in the Materials and Methods.

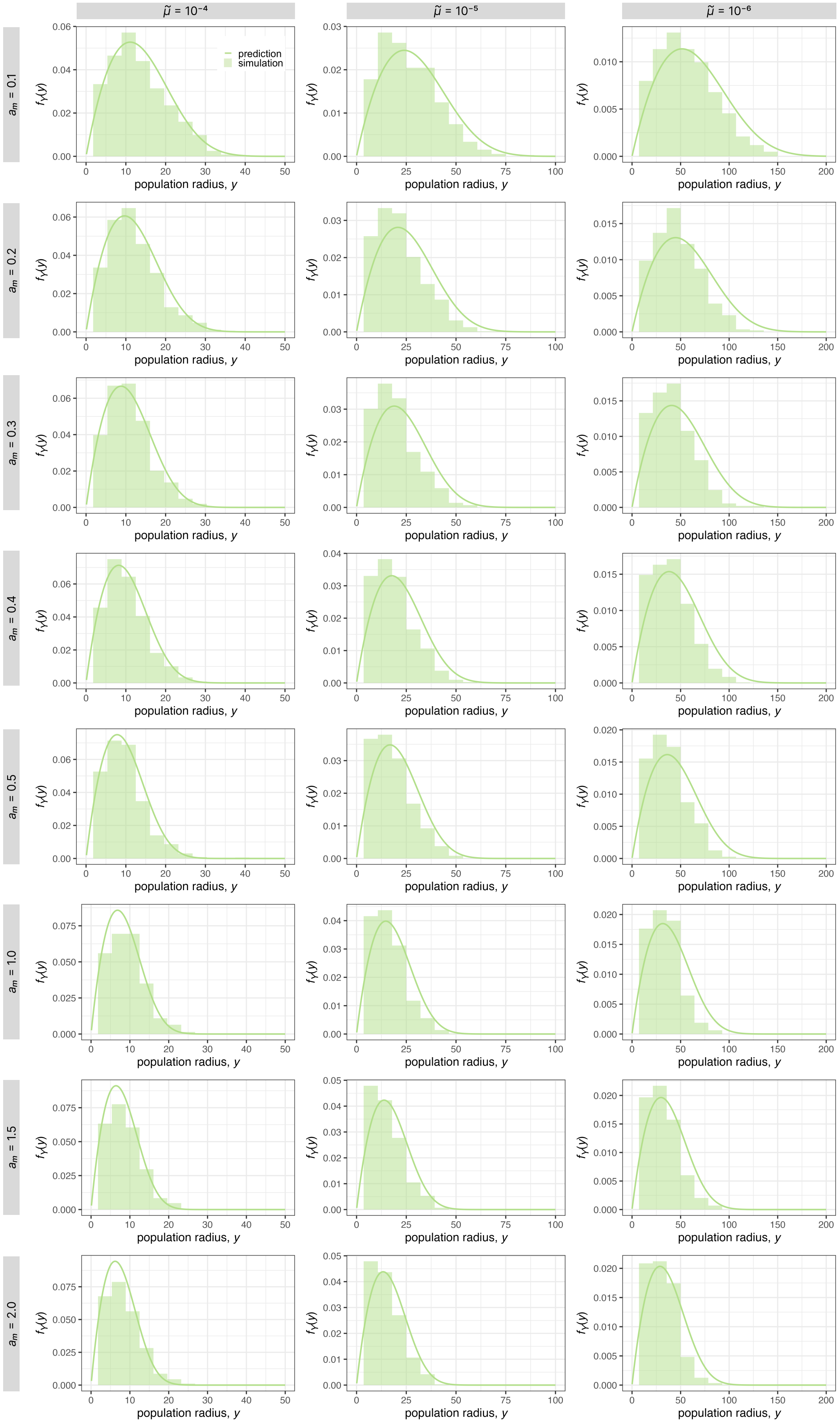


**Fig. S3.** Comparison of simulation results with our analytic predictions of $f_Y$ in 2D for different parameter combinations. We use $m = 0.05$, $K = 16$, $r_{\mathrm{re}} = 0.91$, $r_{\mathrm{wt}} = 1.00$. We vary over $\mu$ and $r_{\mathrm{m}}$ corresponding to different $a_{\mathrm{m}}$. Conversion between simulation and macroscopic model parameters is described in the Materials and Methods.

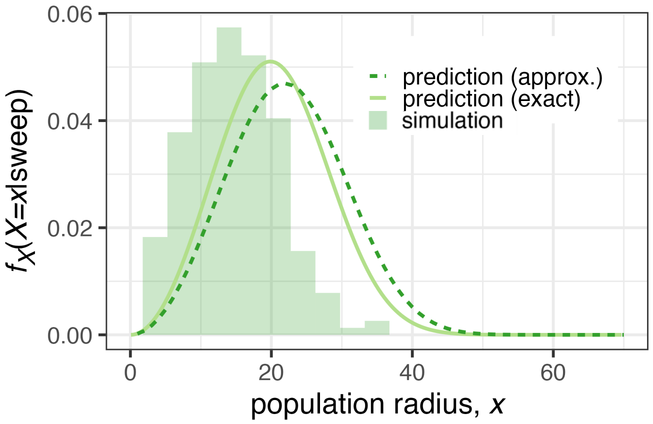


**Fig. S4.** Comparison of simulation results with our analytic predictions of $f_X(x|\text{sweep})$ in 2D. We use $m = 0.05$, $K = 16$, $r_{\text{re}} = 0.91$, $r_{\text{wt}} = 1.00$, $r_{\text{m}} = 1.3$ and $\tilde{\mu} = 10^{-5}$ leading to speeds $c_{\text{wt}} = 0.15$ and $c_{\text{m}} = 0.31$ and survival probability $\rho = 0.23$ (see Materials and Methods).

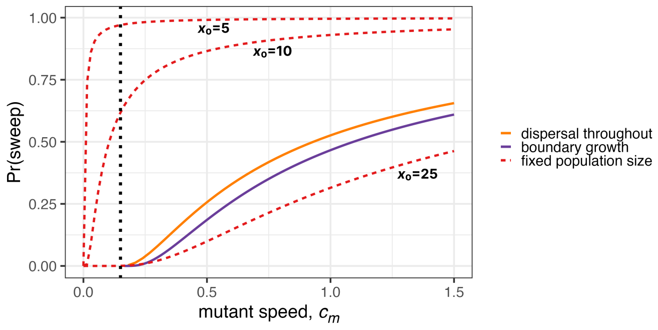


**Fig. S5.** Comparison of sweep probabilities between different growth models in 3D. We show the sweep probability over the mutant speed for models with proliferation restricted to the boundary (solid purple line), population size fixed at radius $x_0$ (dashed red lines), and our main model in which mutants can expand within the expanding wildtype population (solid orange line). In the two range expansion models, the wildtype speed is set to $c_{\text{wt}} = 0.15$ (dotted black line).

 

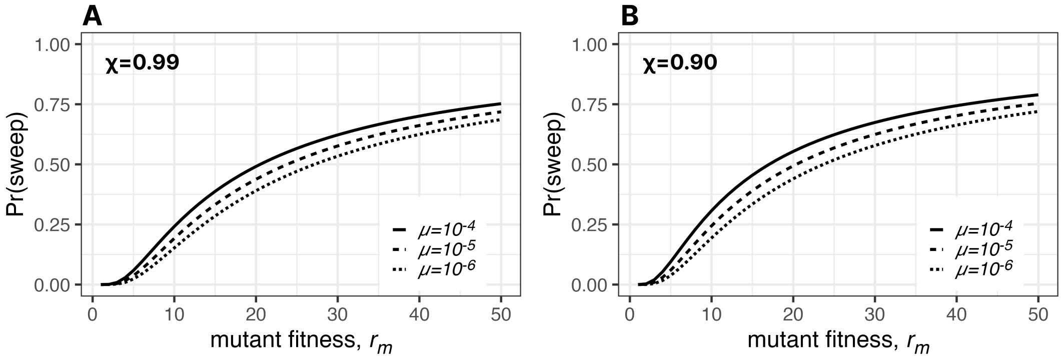


**Fig. S6.** Sweep probability in exponentially growing populations. The wildtype growth rate is set to $r_{\text{wt}} = 1.0$ and we vary over the mutation rate $\mu$ that is conditioned on survival of drift. We say a sweep has occurred once the mutant has reached frequency $\chi$, which we set to $\chi = 0.99$ in **A** and to $\chi = 0.90$ in **B**.

| | 1D | 2D | 3D |
|---|---|---|---|
| $f_X(x)$ | $\frac{2x}{\theta_{1D}^2} e^{-\frac{x^2}{\theta_{1D}^2}}$ | $\frac{3x^2}{\theta_{2D}^3} e^{-\frac{x^3}{\theta_{2D}^3}}$ | $\frac{4x^3}{\theta^4} e^{-\frac{x^4}{\theta^4}}$ |
| $f_Y(y \mid X = x)$ | $\frac{1}{x} 1\{y \leq x\}$ | $\frac{2y}{x^2} 1\{y \leq x\}$ | $\frac{3y^2}{x^3} 1\{y \leq x\}$ |
| $f_Y(y)$ | $\frac{1}{\theta_{1D}} \Gamma\left(\frac{1}{2}, \frac{y^2}{\theta_{1D}^2}\right)$ | $\frac{2y}{\theta_{2D}^2} \Gamma\left(\frac{1}{3}, \frac{y^3}{\theta_{2D}^3}\right)$ | $\frac{3y^2}{\theta^3} \Gamma\left(\frac{1}{4}, \frac{y^4}{\theta^4}\right)$ |
| $\Pr(\text{sweep} \mid X = x, Y = y)$ | $e^{-\frac{x^2+y^2}{\alpha_{1D}^2}}$ | numerical evaluation | $e^{-\mu\left(Ax^4+By^4+Cx^2y^2\right)}$ |
| $\Pr(\text{sweep} \mid X = x)$ | $\frac{\sqrt{\pi}\alpha_{1D}}{2x} e^{-\left(\frac{x}{\alpha_{1D}}\right)^2} \operatorname{erf}\left(\frac{x}{\alpha_{1D}}\right)$ | numerical evaluation | numerical evaluation |
| $\Pr(\text{sweep} \mid X = x, Y = 0)$ | $e^{-\left(\frac{x}{\alpha_{1D}}\right)^2}$ | $e^{-\left(\frac{x}{\alpha_{2D}}\right)^3}$ | $e^{-\left(\frac{x}{\alpha}\right)^4}$ |
| Pr(sweep) [approx.] | $\frac{c_{\text{m}} - c_{\text{wt}}}{c_{\text{m}}}$ | $\left(\frac{c_{\text{m}} - c_{\text{wt}}}{c_{\text{m}}}\right)^2$ | $\left(\frac{c_{\text{m}} - c_{\text{wt}}}{c_{\text{m}}}\right)^3$ |
| Pr(sweep) [exact] | $\frac{\beta'}{\sqrt{1+\beta'}} \cot^{-1}\left(\sqrt{1+\beta'}\right)$ | numerical evaluation | numerical evaluation |
| $f_X(X = x \mid \text{sweep})$ [approx.] | $\frac{2x}{\theta_{1D}^2\beta} e^{-\frac{x^2}{\theta_{1D}^2\beta}}$ | $\frac{3x^2}{\theta_{2D}^3\beta^2} e^{-\frac{x^3}{\theta_{2D}^3\beta^2}}$ | $\frac{4x^3}{\theta^4\beta^3} e^{-\frac{x^4}{\theta^4\beta^3}}$ |
| $f_X(X = x \mid \text{sweep})$ [exact] | $\frac{\Pr(\text{sweep} \mid X=x) f_X(x)}{\Pr(\text{sweep})}$ | numerical evaluation | numerical evaluation |

**Table S1. Summary of analytical results of our main model for 1D, 2D and 3D.**

| Model | Sweep probability | Reference |
|---|---|---|
| boundary growth and cell proliferation throughout the population | $\int_0^\infty \int_0^x e^{-\mu \int_0^\infty N_{\mathrm{wt}}(\tau)\, d\tau\, \frac{3y^2}{x^3} \frac{4x^3 e^{-x^4/\theta^4}}{\theta^4}\, dy\, dx}$ $< \left(\frac{c_{\mathrm{m}} - c_{\mathrm{wt}}}{c_{\mathrm{m}}}\right)^3$ | here |
| boundary growth and cell proliferation restricted to the boundary | $\frac{8+\frac{c_{\mathrm{m}}}{c_{\mathrm{wt}}}}{\frac{c_{\mathrm{m}}}{c_{\mathrm{wt}}}-1} \frac{2}{1+\exp\left(3\pi/(\frac{c_{\mathrm{m}}}{c_{\mathrm{wt}}}-1)\right)}$ | Ref. (7) |
| constant population | $e^{-\frac{\mu \pi x_0^4}{c_{\mathrm{m}}}}$ | Ref. (3, 4) and here |
| exponential growth | $\int_0^\infty e^{-\frac{\mu}{r_{\mathrm{wt}}} e^{rt_1}\left(e^{r_{\mathrm{wt}} t_2(t_1)}-1\right)}$ $\times \mu e^{rt_1} e^{-\frac{\mu}{r} e^{rt_1}}\, dt_1$ | here |

**Table S2. Summary of sweep probabilities for different growth models in 3D.**
